# Mechanical Implications of Cellular Viscoelasticity, Cortex Polarity, Superelasticity, and Cell-Cell Junctions in Curved Tissues

**DOI:** 10.1101/2024.08.01.606202

**Authors:** Amaury Perez-Tirado, Ulla Unkelbach, Tabea A. Oswald, Johannes Rheinlaender, Tilman E. Schäffer, Markus Mukenhirn, Alf Honigmann, Andreas Janshoff

## Abstract

Investigations of the response of curved epithelia derived from MDCK-II cells to external deformation involved indentation-relaxation experiments using colloidal probe microscopy. Notably, hemicysts exhibited lower tissue tension, greater compliance, and increased fluidity compared to cysts. The primary response to deformation turned out to be the in-plane expansion of the basal cortex/membrane of cells. Additionally, drug treatments applied to curved tissue, along with deformation of tailored mutants (such as E-cadherin knockout), revealed that tissue compliance over short time scales is influenced by an interplay of viscoelastic properties in individual cells, their apical-basal polarity, superelasticity of the shell, and excess interfacial area. Meanwhile, tissue resilience predominantly depends on the integrity of cell-cell contacts.

## INTRODUCTION

Epithelial monolayers form physical and chemical barriers and control the permeability between the internal and external environment. These tissues also experience substantial mechanical forces. For instance, the lung experiences enormous intrinsic, short-lived deformations during the breathing cycle,^1^ while epithelia from other organs such as the urothelium sustain long-lasting external deformations originating from pressure gradients.^2, 3^ For optimal tissue resilience, the cells must be mechanically linked to distribute stresses throughout the entire tissue.^4–6^ This integration is accomplished by compliant, viscoelastic cell-cell junctions such as desmosomes,^7^ adherens junction^8, 9^ and also tight junctions^10^ interconnecting elements of the cytoskeleton to generate structures exceeding the length scale of individual cells. Failure to achieve a stable connection can result in tissue fracture, leading to pathological consequences such as hemorrhage and septicemia, whereas the ability to withstand large strain for a prolonged time and to dissipate the absorbed energy protects tissue from fracture.^6^ On a single cell level, excess surface area in conjunction with the contractile actomyosin cortex permits to absorb external stress and to dissipate the associated energy reminiscent of a soft glassy material.^11–15^ However, on tissue length scales, the number of experiments is sparse and our knowledge how cells organized in epithelia respond to lateral, in-plane, forces and their ability to dissipate the elastic energy remains limited.^4–6^

Here, we examine how cysts^16, 17^ and hemicysts (domes) with opposing polarity generated from MDCK-II cells respond to external deformation with micrometer-sized colloidal probes on short time scales of seconds. We found that the compliance of the tissue and its stress relaxation largely mirror the viscoelastic response of single cells governed by the mechanics of the actin cortex and recruitment of excess surface area to alleviate stress. Cell-cell contacts are mainly responsible for the stability of tissue and contribute to the compliance in a nonlinear fashion depending on the number of bonds between cells and the excess interfacial area.

## RESULTS

### Theoretical Model

We assume that curved epithelia subject to external deformation can be described by the Helfrich-Canham Hamiltonian *𝔉* with appropriate boundary conditions:

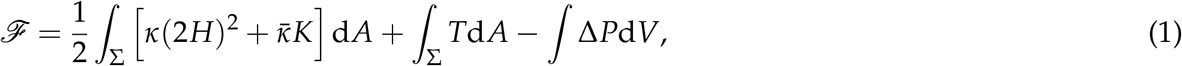

Σ represents the epithelial surface, Δ*P* is the external pressure difference across the epithelia, *H* is the mean curvature, and *K* is the Gaussian curvature. The constants *κ* and 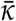 are the splay and saddle splay moduli, respectively. *T* is the Lagrange multiplier to maintain constant area of the epithelia serving as the tissue tension. The shape equation is from standard variational calculus reads:^18^

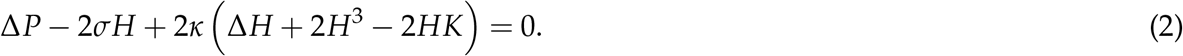

Neglecting the bending contribution (see SI and Fig. SI3), this Euler-Lagrange equation simplifies to the well-known Young-Laplace equation:

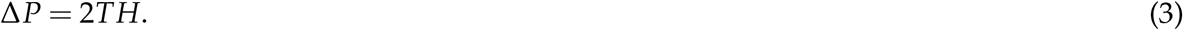

Assuming that the tissue tension *T* is isotropic and depends on area dilatation (Δ*A* = *A*_ind_ − *A*_0_) to first-order, we can write:

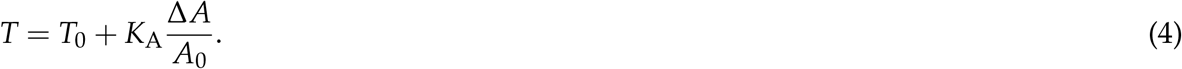

The pre-stress (tissue tension) *T*_0_ of the (hemi)cyst captures the tension of the epithelia prior to deformation due to the existence of Laplace pressure (Δ*P* = 2*T*_0_/*R* for cysts). Below we will see how this pre-stress is relaxed by the thinning of the cells when being dilated. *A*_0_ is the initial area of the (hemi)cyst prior to deformation and *K*_A_ represents the area compressibility modulus. This area compressibility modulus relates to the bulk Young’s modulus of the shell by *E*_Y_ = *K*_A_*h*, with *h* the thickness of the cell layer, i.e., the height of the cells forming the tissue. The shape of the deformed epithelia is a minimal area surface of constant mean curvature parameterized as shown in Fig. SI1/2. Viscoelasticity of cysts is included through the time dependent area compressibility modulus of the shell *K*_A_(*t*) applying the elastic-viscoelastic correspondence principle (see Materials and Methods).^19^

### Preparation and characterization of curved epithelia

MDCK-II cells serve as a representative cell line displaying characteristic features of kidney tubule epithelial cells.^20, 21^ Hemicysts obtained from MDCK-II monolayers maintain the traditional apical-basal polarity, with the apical domain facing towards the media, while the basal domain remains oriented towards the lumen and the surface (Fig. 1A/B). The confocal images reveal that the cells forming the hemicyst are slightly thinner compared to those in the monolayer reminiscent of pre-confluent cells. Conversely, cysts exhibit reversed polarity, with the apical side oriented towards the lumen and the basolateral membrane/cortex facing towards the extracellular matrix (Fig. 1E/F). The topography was further scrutinized using non-invasive, label-free scanning ion conductance microscopy (SICM) (Fig. 1E-H).^22^ Cell-cell borders are only visible in SICM-images of hemicysts, while the surface of cysts shows no individual cells. The apical surface of cells from hemicysts display microvilli visible as rough topographic structures (Fig. 1C-D), while the basal surface of cysts is featureless and flat (Fig. 1G-H) likely due to a thin shell of basal ECM deposited by the cells.

**Figure 1.**
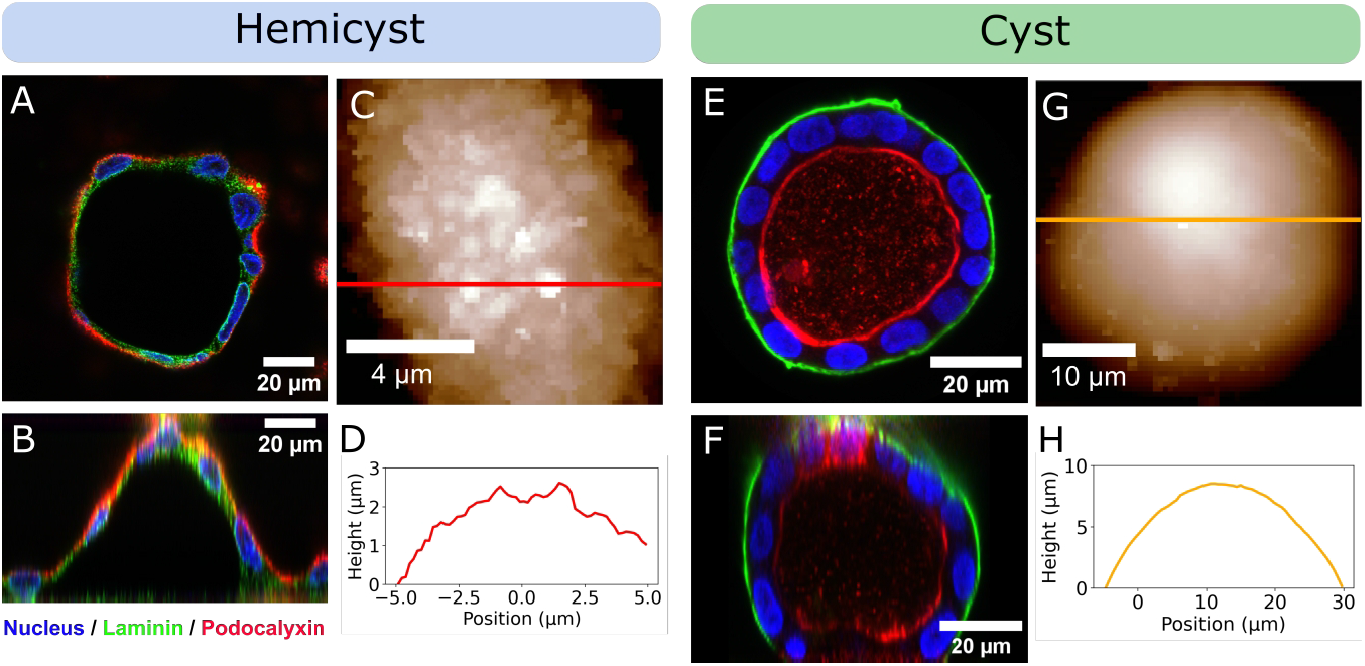
Architecture and topography of cysts and hemicysts. A) Hemicyst of MDCK-II featuring a circular, cell-free space at its center. Nuclei are stained with DAPI (blue, 405 nm), the basal domain (anti-laminin) is shown in green (488 nm), and the apical domain (anti-podocalyxin, GP-135, 546 nm) is depicted in red (). B) Orthogonal view with respect to (A). C) SICM-image of a hemicyst (apical membrane), with the profile marked in red (D). E) Confocal image of a MDCK II cyst captured at its midpoint (staining as in (A). F) Orthogonal view. G) SICM-image of the cyst’s surface (basolateral membrane), with the marked profile shown in (H).

Fig. 2A-H show confocal images of ZO-1 (immunostaining) confirming the described apical-basal polarity of hemicysts and cysts. We also examined the shape of the cells using Cellpose-based segmentation of membrane stained cells (CellMask™, Deep red).^23^ We infer from the projected cell area *A*, the perimeter *P* (Fig. 2I-N). In contrast to hemicysts, the cells forming cysts are forming a compact tissue, typical for a highly jammed/packed state (Fig. 2I-N).

**Figure 2.**
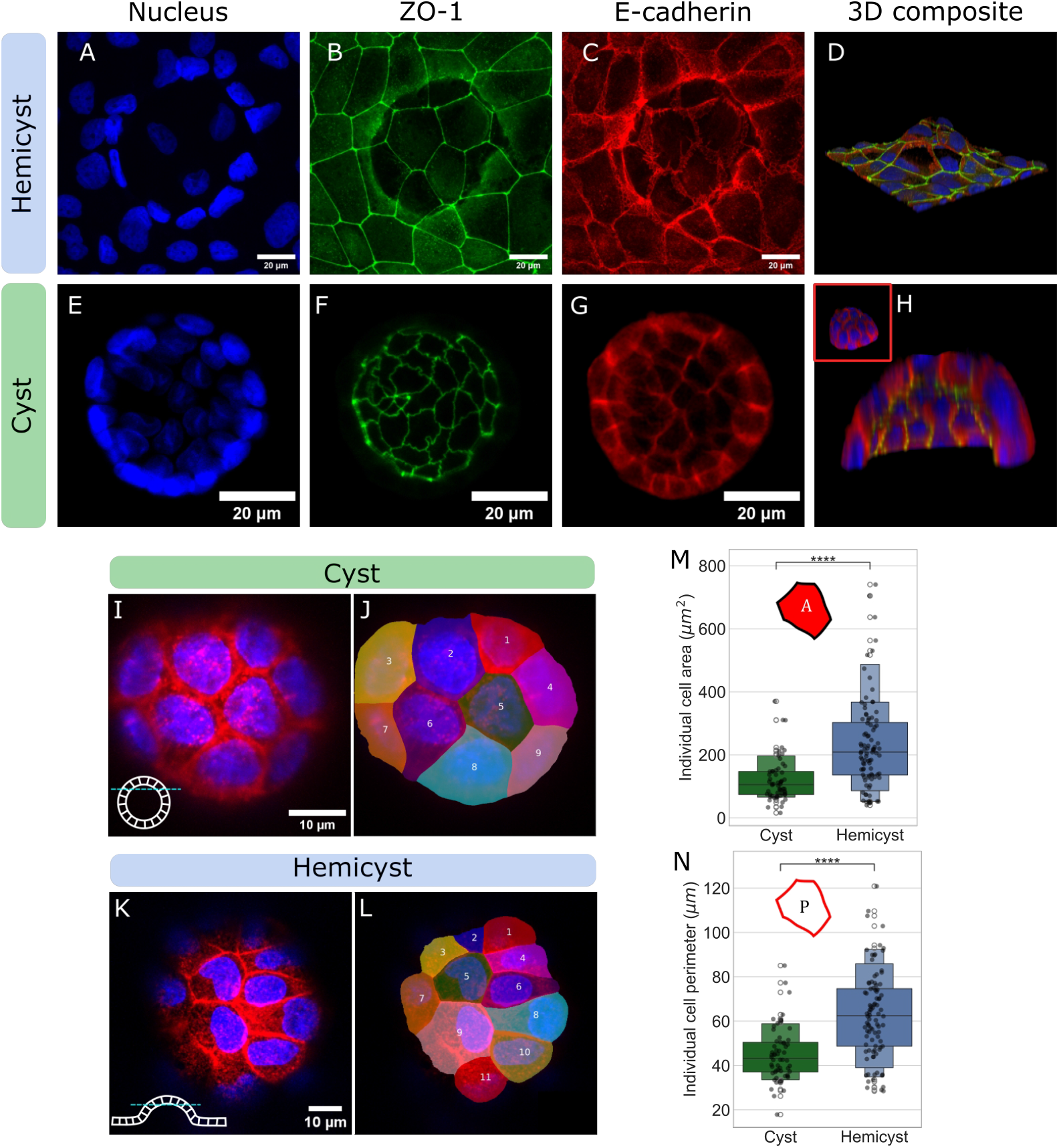
Cell-cell contacts of curved epithelia. Projection of z-stack of confocal images showing nuclei (DAPI, 405 nm), tight junctions (FITC-conjugated Anti-ZO-1, 488 nm), and adherens junctions (Anti-E-cadherin, 546 nm) for hemicysts (A-C) and cysts (E-G). D/H) 3D composite of hemicysts (D) and cysts (H). Cell shape is inferred from CLSM images (nuclei in blue using Hoechst, 404 nm, membrane in red using CellMask™, Deep red, 647 nm) of the top of the structures using the segmentation algorithm Cellpose for cells forming cysts (I-J) and hemicysts (K-L).^23^ Cysts display more compact cells in terms of area (M) and perimeter (N).

### Validation and prerequisites of the tension-based model

The mechanical properties of curved epithelial structures are examined using spherical indenters (*R* = 20 µm) allowing us to approximate the multicellular structures as a continuum. Fig. 3A/B and Fig. SI2 illustrate the experimental setup and the parametrization of the deformed shell. Fig. 3C/D show the average of 396 and 122 indentation-relaxation curves obtained from cysts (3C) and hemicysts (3D), respectively. Cysts, subjected to the same maximum force of approximately 20 nN (1 µm/s), exhibit greater stiffness compared to hemicysts. After indentation, we observe a significant force relaxation indicating viscoelastic properties of the tissue. These properties are further quantified by employing a tension-based model (see SI and Materials and Methods) to describe the data. In the following, we experimentally validated the fundamental assumptions of the model (equations (6,7) and Fig. 3E-G):

**Figure 3.**
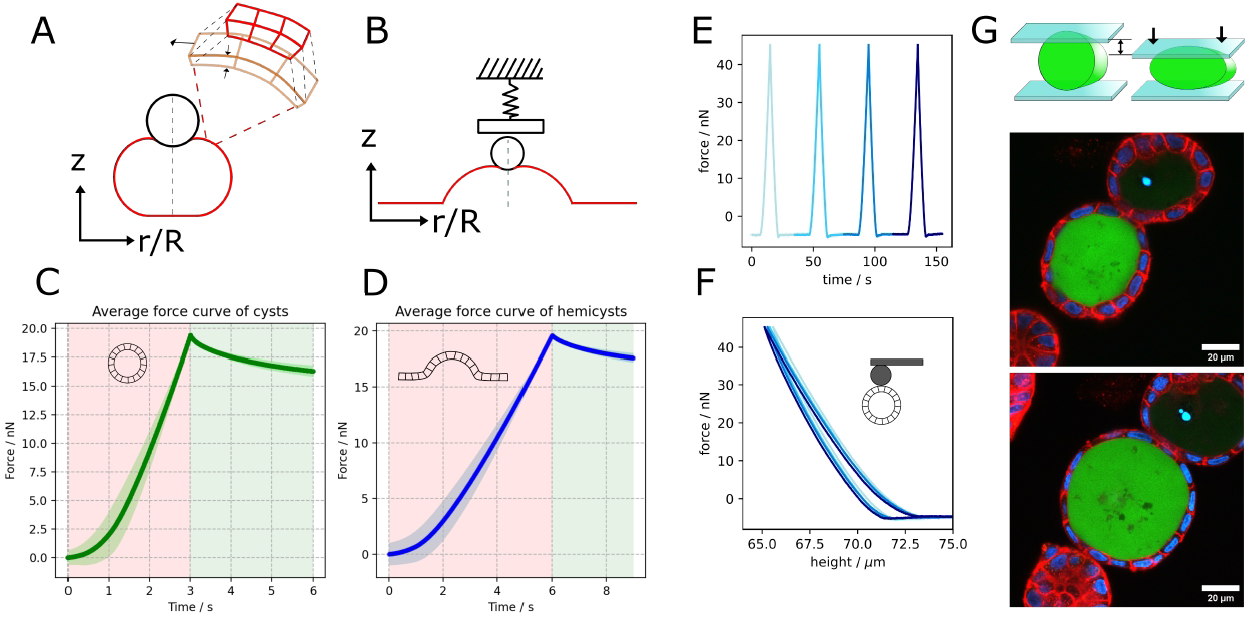
Mechanical probing of cysts and hemicysts. A/B) Scheme of the experimental setup for acquiring indentation-relaxation curves on cysts (A) and hemicysts (B) including the schematic portrayal of superelasticity leading to thinning of the cells in response to area dilatation (inset in (A)). C/D) Averaged indentation-relaxation curves obtained from cysts (C, *n* = 396, green) and hemicysts (D, *n* = 122, blue) deformed with a spherical indenter. Hemicysts are probed apically and cysts are deformed from the basolateral membrane/cortex. E/F) Repeated acquisition of indentation retraction curves on the identical cyst reveals no memory effect, indicating the absence of fluid loss from the lumen at this time scale. G) Compression of cysts between two cover slips shows superelastic thinning of the cell monolayer in response to pressure increase (A-E). The nuclei are stained with DAPI (blue, 405 nm), the cell membrane with CellMask™ (red, 649 nm) and the lumen with the fluorescent labeled mucin (Dendra2-CD164, 488 nm). No efflux of Dendra2-CD164 was detected during compression of the cysts.

i. *Volume conservation of cells and lumen:*Volume conservation of cells subject to deformation has been established in prior research^24^ and can also be inferred from imaging of cysts in the presence or absence of Laplace pressure (Fig. 7). Conservation of the enclosed lumen of cysts and hemicysts is validated by: 1) Conducting consecutive indentation-retraction experiments on the same cyst (Fig. 3E/F), 2) introducing a dye into the lumen and monitoring its efflux during compression with a cover slip (Fig. 3G), and 3) employing different load velocities that should alter the force response if volume conservation was violated (Fig. SI8). We found that taking consecutive force curves (approach-retraction) does not result in visible memory effects indicative of volume loss such as pseudo-softening (Fig. 3E/F). Furthermore, efflux of the endogenously produced dye sialomucin (Dendra2-CD164) from the lumen was not monitored after compression of cysts between two glass slides (Fig. 3G). Finally, we did not notice variation of mechanical parameters after changing the loading velocity by one order of magnitude (Fig. SI8).
ii. *Bending contribution:* bending contributes to the Hamiltonian only when the curvature of the structure is large compared to the shell’s thickness. Figure SI3 illustrates how stretching, pre-stress, and bending each contribute to the restoring force during deformation. At low areal strain, pre-stress is the dominant factor over bending. However, at large strains, stretching becomes more significant than both pre-stress and bending. We used two different indenter geometries (parallel plate and spherical indenter) and probed cysts of various sizes resulting in unaltered tissue tension *T*_0_ (Fig. SI9, Fig. 4).
iii. *Geometry of curved epithelia:* Fig. 1 confirms that cysts are spherical and hemicysts can be described as domes with a contact angle of about 80^°^.
iv. *Linear viscoelastic regime:* As depicted in Fig. SI10, the relationship between area dilatation and indentation depth confirms that the increase in area is sufficiently small (< 1%) to assume tension increases proportionally to *α* = Δ*A*/*A*_0_.

**Figure 4.**
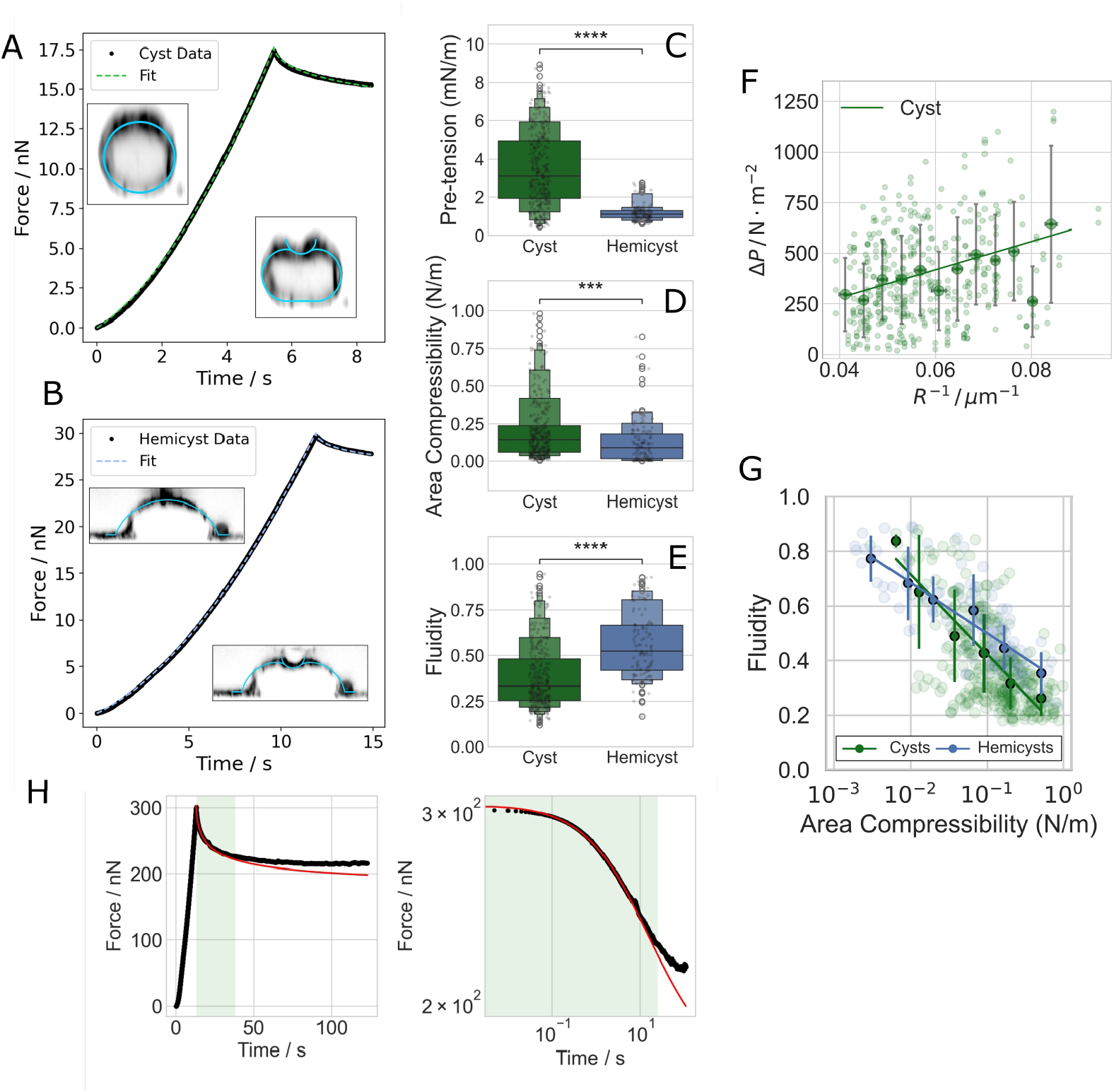
Viscoelasticity of curved epithelia. A) Fitting of the tension-based model (green dotted line, equations (6, 7)) to an indentation-relaxation curve (black dots) obtained from probing a cyst (*R*_c_ = 22 µm) at a maximal indentation depth of 5.34 µm providing *T*_0_ = 1.85 mN/m, 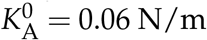 and *β* = 0.34. The images (inverted colors) show the z-projection of the middle slices taken from *xz*-cuts of CLSM *z*-stacks, before (top) and after (bottom) indentation (200 nN), superimposed with the corresponding shape obtained from the tension-based model (blue line). B) Indentation-relaxation curve obtained from a hemicyst subject to fitting of the tension-based model with adapted geometry (blue dotted line corresponds to the fit, the black dots are the experimental data). The base radius of the hemicyst was approximately *R*_hc_ = 41 µm and the maximal indentation depth 12 µm leading to *T*_0_ = 0.53 mN/m, 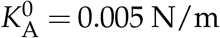 and *β* = 0.4. The images depict the shape of a hemicyst before (top) and after (bottom) indentation (200 nN), overlaid with the corresponding model shape (blue line). C-E) Pre-tension (*T*_0_), area compressibility modulus 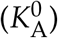 and fluidity *β* of cysts (*n* = 346) and hemicysts (*n* = 115). F) Laplace pressure of cysts computed from tension values as a function of curvature. The slope of the linear regression (black line) provides an average tension 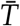 of 3.5 *±* 0.3 mN/m. G) Fluidity (*β*) as a function of stiffness represented by 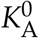. H) Indentation-relaxation curve of a cysts deformed with a spherical indenter (*R* = 20 µm) (left: linear plot, right: semi-logarithmic plot).

### Viscoelastic properties of hemicysts and cysts

Fig. 4A/B shows two exemplary fits (dotted lines) of equations (6, 7) to indentation-relaxation curves acquired from deforming cysts (Fig. 4A) and hemicysts (Fig. 4B) with a spherical indenter. These fits aptly describe the data, and the scaling analysis (Fig. SI11) substantiates this observation. Three parameters, pre-stress *T*_0_, area compressibility modulus (scaling factor) 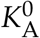 and the fluidity *β* are obtained and shown in Fig. 4C-E. Fig. SI12 shows how the viscoelastic parameters impact indentation-relaxation curves. Relaxation was limited to few seconds, since we found that the power law describes stress relaxation only up to 25 s. After 25 s, relaxation saturates and a stationary state is reached (Fig. 4H). During this time cell movement, shape changes or cell divisions were not detected.

Further confirming the applicability of the tension model, the final shapes of the deformed epithelial structures and the ones predicted from the model were compared showing convincing congruence (insets in Fig. 4A/B).

#### Pre-stress of cysts and hemicysts

Cysts exhibited a significantly higher mean pre-stress *T*_0_ of 3.4 *±* 1.9 mN/m (mean *±* standard deviation) than hemicysts (*T*_0_ = 1.2 *±* 0.4 mN/m) (Fig. 4C). According to their larger curvature, cysts also experience significantly greater Laplace pressure than hemicysts (< 50 Pa). The consistency of the tension model is demonstrated for cysts in Fig. 4F. As expected for a tension-based response, the Laplace pressure varies linearly with curvature consistent with size-invariant description of the deformation. The slope (black line in Fig. 4F) corresponds to the average tension 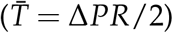 and is essentially identical to the mean tension obtained from individual indentation relaxation curves shown in Fig. 4C.

#### Intrinsic viscoelastic parameters

The area compressibility modulus 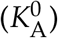 represents the stiffness of the curved epithelial shell and becomes dominant at larger areal strain (Fig. SI12). It reports on the cell-cell contacts including excess interfacial area as well as the mechanical properties of individual cells. Here again, hemicysts show lower values of 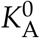 in comparison to cysts (Fig 4D). The bulk elastic modulus of the curved epithelia can be readily obtained from the area compressibility modulus (scaling factor) and the thickness of the shell by *E*_Y_ = *K*_A_/*h*.

From optical microscopy images we obtained an average thickness of the curved epithelia of *h* = 6.5 *±* 1.1 µm for cysts and *h* = 7.4 *±* 1.2 µm for hemicysts, providing us with *E*_Y_ = 26.7 *±* 3.2 kPa for cysts and *E*_Y_ = 17.2 *±* 4.1 kPa for hemicysts. Hemicysts also exhibit greater fluidity compared to cysts, as indicated by the power law exponent *β* (Fig. 4E). Hence, cysts lean towards a more solid-like behavior. Interestingly, suspended monolayers show even lower mean *β*-values of 0.15 *±* 0.03.^4^ Fig. 4G demonstrates that fluidity (*β*) and stiffness 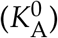 are not independent as also found for individual cells.^12, 13, 15, 25^ Furthermore, the limit of fluidity is found to be approximately 0.2 as also reported for isolated cortices.^26^

### Superelasticity and stress stiffening

It is anticipated that curved epithelia will exhibit superelastic behavior in response to changes in external or internal pressure.^27^ Uniform Laplace pressure causes the cells and the nuclei to thin and thereby reducing the lateral stress in the monolayer. Fig. 3G and Fig. 5A-C which illustrate how cells composing cysts flatten in reaction to the presence of Laplace pressure. Fig. 5 shows how the sudden loss of Laplace pressure leads to decrease in area expansion of *α* ≈ 100 %. Concomitantly, the cell layer thickness (Δ*h* ≈ 100 %) as the tension vanishes. The volume of the individual cells is conserved during compression and the excess area stored in the apical side of the cells pointing to the lumen is recruited during pressure buildup. Indentation with moderate forces well below 100 nN generate only moderate area dilatation on the order of 1%. Hence the impact of superelasticity during indentation and relaxation will be small and can be lumped into a reduced apparent pre-stress (see SI).

**Figure 5.**
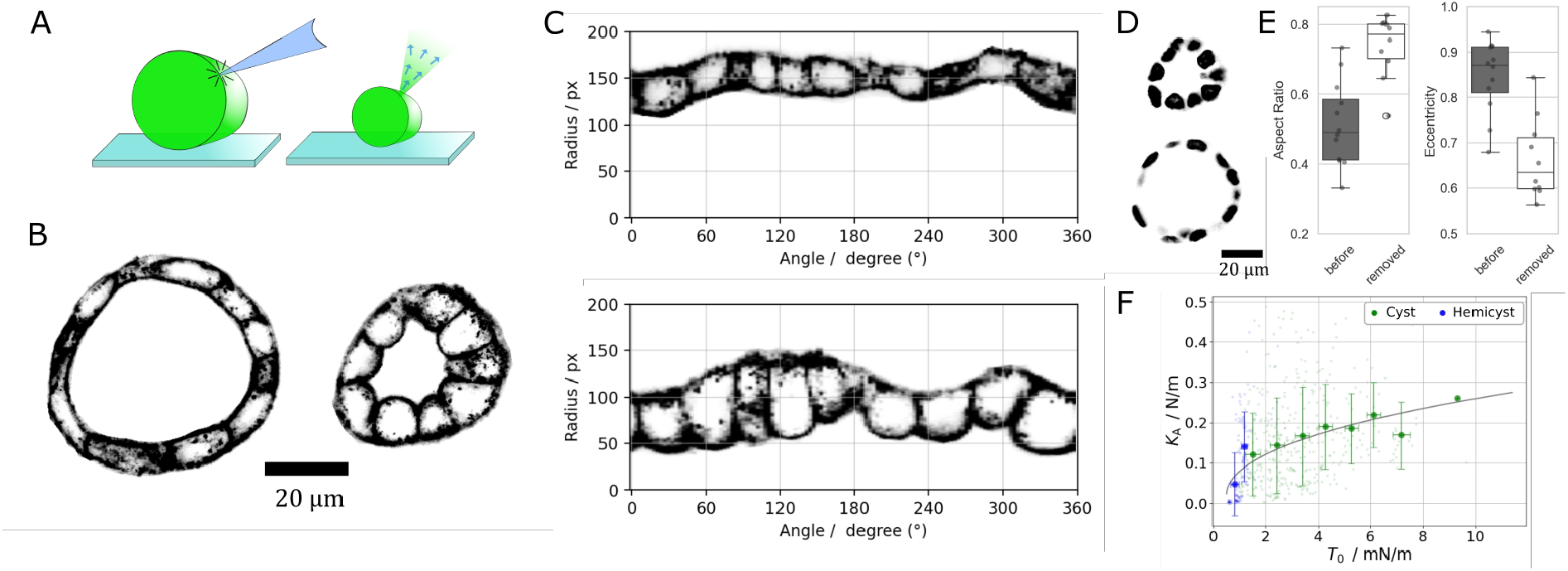
Superelasticity of curved epithelia. A) Scheme illustrating the permeation of cysts with a micropipette. B/C) Volume conservation forces the cells back to their original shape by expanding apically within the lumen resulting in an increased cell height and decreased circumferential dilation (top of C): before leakage; bottom of C): after leakage). The nuclei are stained with DAPI (blue 405 nm), the membrane with CellMask™ (red 649 nm) and the lumen with the fluorescent mucin (Dendra2-CD164). The images were color-inverted and shown in black and white. D) Nuclei image extracted and color-inverted for analysis. E) Nuclear shape parameters of the pressurized and deflated cysts. F) Area compressibility modulus (scaling factor) 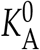 as a function of pre-stress *T*_0_ that reports on the initial areal strain. Blue dots refer to hemicysts and green dots to cysts. The black line denotes a plot of equation (11) with 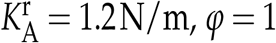 and 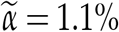.

However, superelasticity might be also responsible for the observed nonlinear stress-stiffening in epithelia. Fig. 5F shows how the stiffness of the epithelia 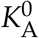 derived from hemicysts (blue) and cysts (green) changes as a function of the measured pre-stress *T*_0_. Assuming a linear relationship between this pre-stress and area dilatation *α*_0_ we can describe the data with equations (10,11).

The apparent stiffness (scaling factor) 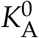 of hemicysts increases strongly with pre-tension (*T*_0_), while the apparent stiffness of cysts is almost constant. The intrinsic stiffness derived from the fit is approximately substantially higher 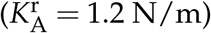 than the measured one.

### Single Cell Mechanics

One key question to be answered is to which extent the response of the curved epithelia is influenced by the viscoelastic and morphological properties of the individual cells that form the supracellular structure (Fig. 6). Employing conventional nanometer-sized AFM-probes, we centrally indented the cells of curved epithelia and fitted the indentation-relaxation curve to a recently developed model, giving access to the cortical tension, the cortex’s area compressibility modulus, and its fluidity.^13–15^ Crucially, we encounter two distinct orientations of the cells: in hemicysts, the more bulged apical side faces towards the probe, whereas we target the rather flat basolateral membrane/cortex of the cells in cysts (Fig. 1). As the model requires the geometry of the unperturbed cells and extracellular matrix forming the curved epithelia we estimated the average cap dimensions from topographical imaging (Fig. 1C/G and Fig. SI13). The fits impeccably describe the experimental data comprising both indentation and relaxation (Fig. 6A/B). Notably, the apical side of cells from hemicysts and the basal side of cells from cysts exhibited basically identical cortical pre-stress but almost an order of magnitude higher pre-stress than confluent MDCK-II cells grown on regular culture dishes (Fig. 6C).^15^ The area compressibility modulus (scaling factor) was found to be considerably smaller for hemicyst cells compared to cells organized in cysts (Fig. 6D). As recently demonstrated, the stiffness of cells mirrors both the cortex architecture and the excess surface area.^14, 28^ Hence, we infer, supported by confocal and SICM-imaging (Fig. 1), that this excess area is apically larger. Concomitantly, the fluidity was found to be substantially increased in cells from hemicysts compared to cells forming cysts (Fig. 6E). This was expected from the known interdependence of stiffness and fluidity proposing an universal viscoelastic trajectory.^12, 15, 29^ The intrinsically higher softness/fluidity of cells from hemicysts goes hand in hand with the polarity of cells in which the basolateral membrane/cortex is essentially stiffer than the apical side.^30^ In addition, cells forming cysts are in a more mature state displaying a compact morphology indicative of jammed cells in late stage confluent cell monolayers (Fig. 1, SI).^28^ For a quantitative analysis of cell size, we refer to Fig. 2I-P) revealing that cells within cysts are showing a higher degree of apical-basal polarity and exhibit greater compactness. They occupy less area compared to cells in hemicysts, which conversely share essentially identical morphology with adjacent cells attached to the substrate (Fig. SI14). In Fig. SI15 we show representative scanning electron microscopy images of a large number of collapsed hemicysts confirming that cell morphology of cell from hemicysts and the adjacent cells of the adherent monolayer are very similar.

**Figure 6.**
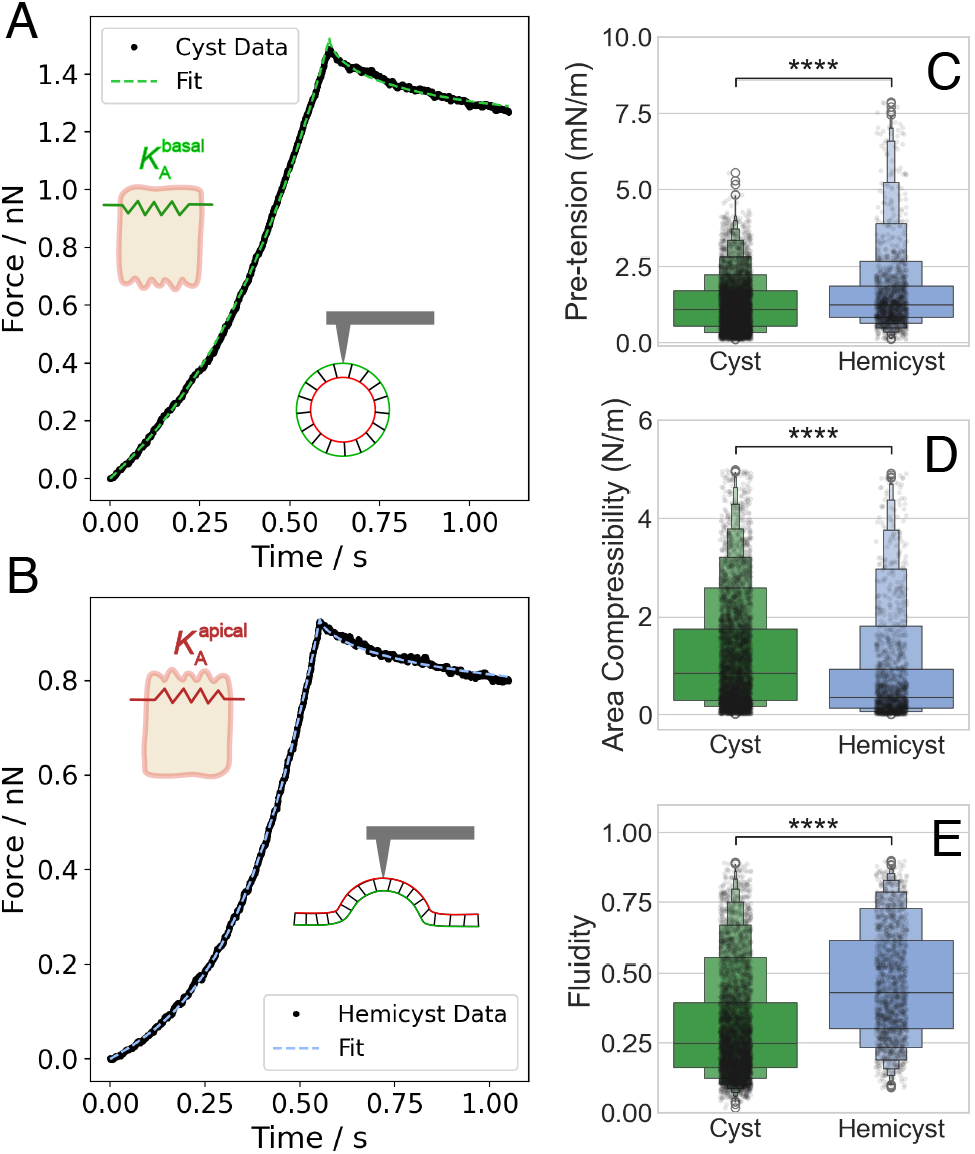
Single cell mechanics. Curve fitting of the indentation-relaxation data obtained from individual cells with a sharp indenter using a tension-based model to describe the response of the cells.^15^ A/B) The graphs depict exemplary fits of indentation-relaxation data obtained from cells forming cysts (A, green dotted line) and hemicysts (B, blue dotted line). A small schematic indicating the polarity that face the indenter and the base is added, apical in red and basal in green. The base radius of the cells was assumed to be 7.5 µm as obtained from imaging. Single cells of a cyst were indented to a maximal depth of 0.61 µm. The mechanical parameters obtained from the fit are *T*_0_ = 1.8 mN/m, 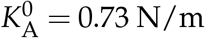 and *β* = 0.28 for cells from the cyst and *T*_0_ = 0.7 mN/m, 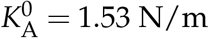 and *β* = 0.12 for cells from the hemicyst. C-E) Viscoelastic parameters derived from cells organized in cysts (green, *n* = 4378) and hemicysts (blue, *n* = 1376).

**Figure 7.**
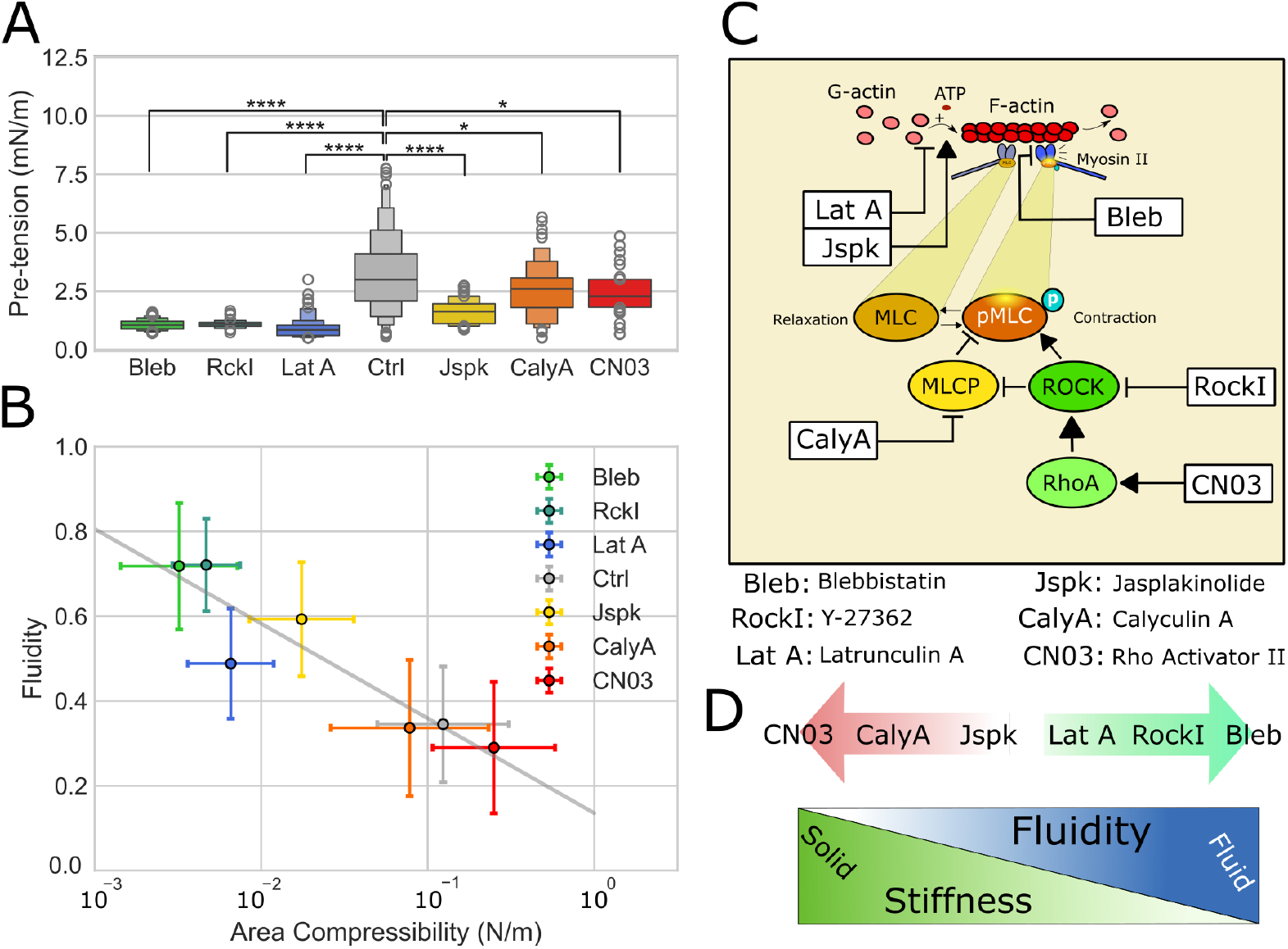
Effects of cytoskeletal drugs on cyst mechanics. A) Pre-tension *T*_0_ of WT-MDCK-II (Wild type) cyst exposed to different cytoskeletal drugs. B) Fluidity (*β*) as a function of area compressibility modulus 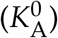 for the cysts exposed to different drugs. The cytoskeletal drugs used were: latrunculin A (Lat A), blebbistatin (Bleb), Rho Activator II (CN03), rock inhibitor Y27362 (RckI), jasplakinolide (Jspk), calyculin A (CalyA). C) Diagram displaying the known targets of the selected drugs. D) Illustration of the effects of drug treatment on fluidity and compressibility.

### Impact of cortex architecture and cellular contractility on tissue mechanics

Earlier reports on tissue-scale mechanics indicate that stress relaxation in epithelial monolayers is controlled by the actomyosin cortex within individual cells.^5^ In the context of epithelia, we know that cortical tension balances adhesion between cells. In our study, we investigated the effects of various drug treatments targeting the contractility and integrity of the actomyosin cortex on the cysts’ mechanical properties (Fig. 7). Our results indicate that inhibiting the contractility of the cortex using either blebbistatin or the Rock inhibitor Y27362 significantly decreases tissue tension (Fig. 7A), while concurrently increasing fluidity, as illustrated in Fig. 7B. This transition towards a more fluid behavior, coincides with a reduction in the shell’s stiffness by more than one order of magnitude. Similar trends emerge when latrunculin-A is employed to disrupt the cortex architecture, while the Rho-activator CN03 maintains tissue tension but stiffens and solidifies the treated cysts. Interestingly, calyculin A, known to increase cortical tension in cells, shows no significant impact on cell rheology in our hands, which we attribute to local effects.^4^ Collectively, these findings suggest that cysts exhibit relaxation behavior akin to individual cells, with the presence of a contractile cortex playing a pivotal role in determining the tissue’s viscoelastic properties, especially constituting the tissue tension *T*_0_. This observation is consistent with prior studies, such as the work by Charras and colleagues, who also found that softening of a cell monolayer results from the addition of latrunculin-B.^5^ Unlike in planar monolayer experiments,^5^ we do not observe a two-step relaxation process on short time scales, or at least, we cannot distinguish it from the power law at our time resolution in the millisecond regime. This might be due to our more involved relaxation function (eq. (7)) that better captures stress relaxation on short time scales than a simple exponential. Conversely, we also observed saturation of relaxation ending in a force plateau on longer time scales (Fig. 6H). Consequently, we conclude that force relaxation within the monolayer is controlled by cortex remodeling and actomyosin flow on the time scale of seconds, similar to the mechanism described for individual cells.^13^ Fig. 7B shows that the dependence of fluidity and stiffness (Fig. 4I) continuous also for the different drug treatments.

### Cell-Cell Junctions

The effect of cell-cell junctions on tissue mechanics was evaluated by using EDTA to broadly weaken these junctions, and specifically by utilizing E-cadherin 1 knockout (Cdh1-KO) cells.^31, 32^ Moreover, Fig. 8A-G show that ZO-1 expression and location are unaltered in Cdh1-KO-cysts, while E-cadherin is clearly missing in between the cells.^31^ Examining the size distribution of cysts selected for mechanical analysis showed that Cdh1KO-cysts are generally smaller than WT-cysts. Naturally, larger cysts experience less area dilatation upon indentation compared to smaller cysts at the same load. Therefore, we divided the WT-cysts into two groups: one with smaller diameters to match the size of Cdh1-KO cysts, and another with larger diameters (Fig. 8H).

**Figure 8.**
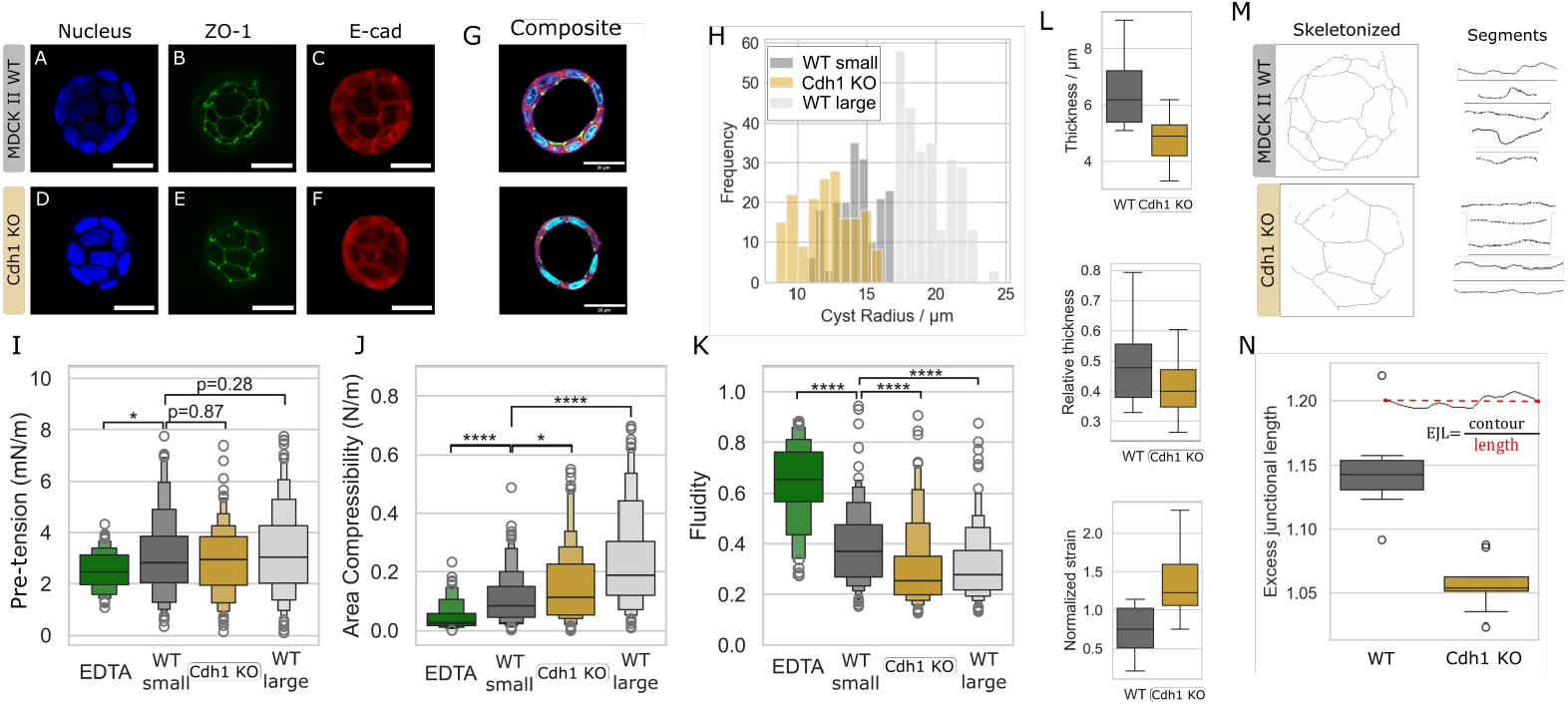
Influence of E-cadherin knockout on cyst mechanics. A-F) Confocal images of E-cadherin-deficient cysts versus WT-MDCK-II cells. Nucleus (A/D, DAPI-staining, blue, 405 nm), ZO-1 (B/E, immunostaining, green, 488 nm), and E-cadherin (C/F, immunostaining, red, 546 nm). G) Composite confocal image (slice) showing that cells forming WT-cysts are less compressed than those in Cdh1KO cysts (E-cadherin 1 knockout). H) Distribution and selection criteria for cyst radii. I-K) Pre-tension, area compressibility, and fluidity of WT-MDCK-II (large and small cysts, see H)), Cdh1-KO, and MDCK-II cells in the presence of 5 mM EDTA. Tension across all cysts is almost identical, while compliance and fluidity of cysts exposed to EDTA are substantially increased. Cells lacking E-cadherin are stiffer and less fluid than WT-cysts of the same size. L) Shape parameters of the Cdh1-KO cells in comparison to WT-cysts. M) Excess apical interfacial area between cells obtained from the ZO-1 contour length normalized to end-to-end distance from node to node. A node occurs where more than two cells meet. Cysts formed by Cdh1-KO cells are more dilated than WT-cells, thus showing less excess interfacial area apically (N).

The mechanical data derived from indentation-relaxation experiments reveal that the pre-stress (*T*_0_) remains consistent in all cysts regardless of the mutation, EDTA administration or diameter (Fig. 8I). Conversely, the stiffness parameter 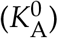 exhibits a marginal increase in Cdh1-KO-cysts, while EDTA-treatment results in a substantial decrease of 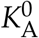 compared to WT-epithelia (Fig. 8J). Similarly, the fluidity parameter *β* is reduced in Cdh1-KO cells, while EDTA-treated cysts show a considerable fluidization relative to WT MDCK-II cells (Fig. 8K).

The finding that cysts lacking E-cadherin show increased stiffness is unexpected, as one might expect that fewer cellular bonds would lead to a softer tissue. We attribute this unexpected finding to the difference in excess interfacial area of the cells forming the epithelia. Fig. 8L shows that cysts formed by Cdh1-KO cells are more compliant apically resulting in a reduced thickness compared to WT-MDCK-II cells at the same Laplace pressure (Fig. 8I/H). This indicates a larger areal strain. Deforming these highly dilated cysts results in larger apparent area compressibility moduli in comparison to the less compliant WT-cysts. The recruitment of excess area is is further supported by the ZO-1 staining shown in Fig. 8M/N. Cdh1-KO-cysts show almost straight lines with no excess junction length in contrast to the zigzag structure found for WT-cysts indicating excess interfacial area and a less stressed strained state.

## DISCUSSION

From a mechanical perspective, the response of epithelial tissue to external deformation involves various length and time scales.^6, 33^ At larger scales, both in length (spanning many cell diameters) and time (hours), the collective viscoelastic properties are primarily influenced by cell rearrangements, cell division and death, along with elastic deformation.^34^ At smaller length scales (micrometers, cell size) and shorter time scales (seconds-minutes), cellular properties comprising rheological properties of the cytoskeleton and cortical contraction prevail.^13, 35–37^ Finally, at the molecular scale (nanometers), intercellular adhesion molecules connect cells with forces in the 10-200 pN range.^38–40^ These molecules are linked to the cytoskeleton, undergo continuous turnover and engage in stochastic bond formation and rupture.^39, 40^

Here, we focused on the mesoscale level, where the sheet’s overall compliance and force relaxation depends on the basolateral and apical cortex of tissue as well as their intercellular connections (Fig. 9). The response to deformation comprises several cell diameters and the time scale of seconds ensuring that no contributions from tissue remodelling are taking place. In the context of stressed tissues, each cell bears a load in the nN-µN regime distributed among the protein complexes forming cell-cell junctions linking neighboring cells.^41^ Therefore, the force scale addressed here ranges from 1 to 200 nN to capture the important stresses. Previous experimental observations revealed a biphasic stress relaxation in tissues, transitioning from fast to slow relaxation, indicative of solid-like behavior at minute timescales. This behavior mirrors findings in indentation experiments, aligning with power-law rheology.^13, 42^ Hence, we compiled both the resilience of the epithelial sheet and its relaxation into a time-dependent area compressibility modulus (Fig. 9). Up to 25 s after the maximal force is reached, stress relaxation is well-described by a power law, while at longer times we find that the force reaches a plateau similar to stress relaxation of a Kelvin-Voigt model (Fig. 4H). Broadly, our tension-based continuum model treats the tissue as a two-dimensional superelastic sheet subject to moderate areal strain. Assuming volume conservation, the resulting shape is therefore adopting a minimal surface. The model not only effectively describes the indentation-relaxation curves obtained from probing of cysts and hemicysts but also accurately predicts the resulting shape of the tissue at maximal deformation (Fig. 4A/B). Importantly, we found that cysts are stiffer and more solid-like than hemicysts, while the pre-stress is basically reflected by the polarity of the cells. Remarkably, all tissue experiments can be mapped onto a master curve, where fluidity and stiffness are intrinsically interconnected. This provides a trajectory along which changes in viscoelasticity are constrained (Fig. 4I and Fig. 8B). We explain the observed trajectory that inversely correlates stiffness with fluidity as a consequence of the conformational energy landscape generated by the cross-linked actomyosin cortex. This landscape contains numerous kinetic traps next to high energy barriers. When the network undergoes elastic deformation, the filaments remain within energy wells. However, if a filament escapes its trap—perhaps due to myosin activity—the stored deformation energy is dissipated as heat.^12^

**Figure 9.**
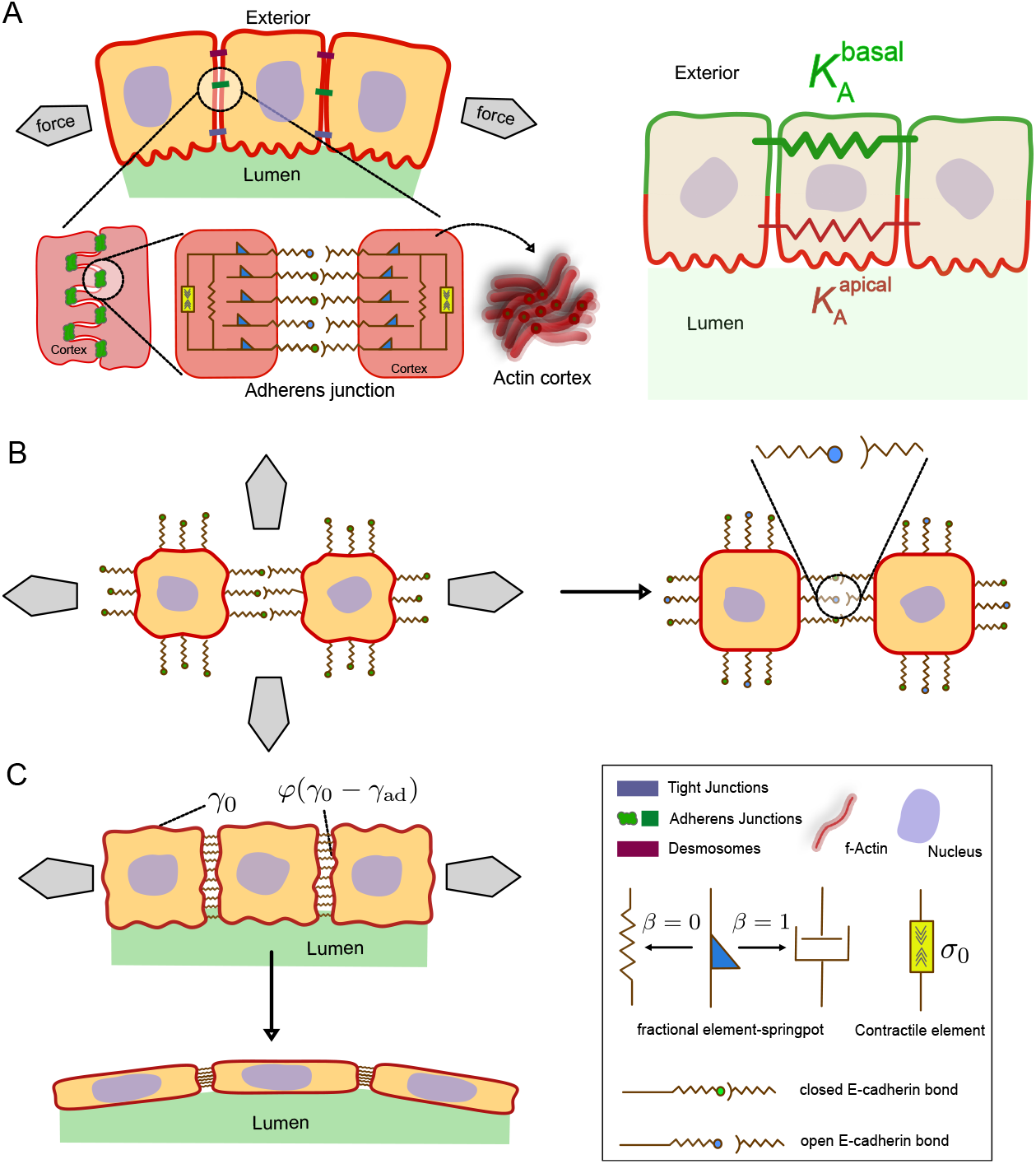
Viscoelasticity contribution of the components in curved epithelial tissue. Scheme illustrating the response of cells arranged in pre-stressed sheets forming curved epithelia like cysts or domes to areal strain. Resilience of the tissue originates from cell-cell-junctions, while the cross-linked actomyosin cortex generates tissue tension and passively acts as a complex fluid represented by fractional viscoelastic elements that dissipate energy. Excess surface area mitigates tension by reducing area dilatation. A) Cross-sectional view of polar curved tissue subject to in-plane tension. When the tissue is stretched by external forces, the resulting areal strain loads cell-cell connections, while the stress dissipates via the cross-linked cortex structure of individual cells. Adherens junctions are modeled by viscoelastic elements consisting of parallel bonds representing homotypic E-cadherin-bonds coupled to the contractile cytoskeleton. Excess interfacial area stored in fold wrinkles and intercalation of the cells is recruited to mitigate the stress acting on the junctions. The apical side pointing to the lumen can recruit more excess area and has been shown to be softer than the basolateral membrane/cortex (right scheme). B) Top view of the stretching process revealing the strain placed on cell-cell connections and the cortex, as well as the reduction of interfacial area in this process showing elongated cell-cell contacts. C) Cross sectional view of the same process illustrating the effect of superelasticity, i.e., thinning of the cells.

Resorting to a purely elastic view, the Young’s moduli of the two-dimensional epithelial sheets (*E*_Y_ = *K*_A_/*h*) align with prior research.^4^ For instance, Harris *et al*. found elastic moduli around 20 kPa, while our data indicates approximately 27 kPa for cysts and 17 kPa for hemicysts. The observed viscoelastic behavior of cysts and hemicysts primarily reflects the properties of the actomyosin cortex emphasising the apical-basal polarity of the cells forming the tissue (Fig. 9A). In cysts, the basolateral membrane/cortex of cells facing outwards is less compliant than the apical side, mostly due to the substantial excess surface area of the apical side mitigating in-plane stresses (Fig. 5).^15^ Hence the primary response to deformation is the in-plane expansion of the basal cortex/membrane of cells. This also explains why bending can be neglected as a major contribution to the Hamiltonian as this cortex is extremely thin, on the order of only 100-200 nm.^11^ In hemicysts, the cell layer is partially compressed before the indenter reaches the basolateral membrane/cortex, which explains the soft response at low indentation depth compared to cysts. Superelasticity further reduces the apparent stiffness measured in force relaxation experiments as additional surface is created at the expense of shell thickness. Eventually, the area compressibility modulus becomes independent of areal strain (Fig. 5).

In cells, the apical actomyosin is linked to the cell’s apical junctions forming structural links to the contractile cortex of adjacent cells.^43^ Hence, these apical junctions attract and organize components necessary for the creation and reinforcement of apical F-actin, as well as for triggering the activity of myosin II enabling the contraction of the cortex.^33, 44^ The junctions provide a viscoelastic link between the cells, which can modulate the rheological properties of the tissue.^8, 9, 45–47^ Hence, perturbations or even disruption of these junctions alter tissue mechanics significantly, highlighting their role in maintaining integrity and creating stiffness.^6, 48^ Cysts treated with low EDTA-concentrations maintained tissue integrity and endure modest areal strain during deformation.^4^ In indentation-relaxation experiments the treated cysts display unaltered tissue tension (*T*_0_) but reduced stiffness paired with higher fluidity compared to untreated WT-cysts. As intercellular bonding is largely impaired by EDTA administration, the loss of cell-cell contacts goes hand in hand with a reduced elastic modulus of the whole shell. The following simplified model illustrates this further. We envision a patch of *n* parallel springs with diameter *D* connected elastically to the cytoskeleton to represent an adherens junction (Fig. 9, Fig. SI16/17). Let *n*(*T*) be the number of closed bonds connecting the cells and forming intact springs. A higher tension *T* leads to bond failure and less springs remain to carry the load. The force acting on the single spring representing the cytoskeleton is 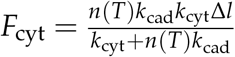, in which Δ*l* is the extension, *k*_cad_ the spring constant of a single cell-cell adhesion molecule such as cadherins, and *k*_cyt_ denotes the spring constant representing the elastic coupling to the actin cortex. The associated elastic modulus arises from Hooke’s law relating stress *σ* to strain (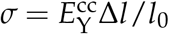, with *l*_0_ the resting length of the serial connection of springs). As the stress can be written as *F*_cyt_/*A*_c_ = *k*_tot_Δ*l*, with the contact area of the patch *A*_c_ = *πD*^2^, we have 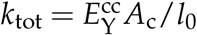. As the number of bonds decreases due to the addition of EDTA, the overall spring constant is 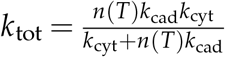 and hence also the effective Young’s modulus of the cell-cell junction 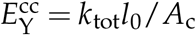 decreases non-linearly with diminishing number of bonds (Fig. SI16/17). Hence, the epithelium’s resilience against stretching and rupture hinges on the massive number of parallel bonds. However, removing a few bonds doesn’t have a substantial impact on the overall Young’s modulus since the load is distributed non-linearly across many bonds.^49^ This explains why cysts depleted only from E-cadherin show no softening since other cadherins such as K-cadherin step in to compensate for the loss. Conversely, reducing calcium levels by adding EDTA decreases the number of closed bonds more drastically, thus resulting in pronounced tissue softening (Fig. SI16/17).

We observed that E-cadherin-depleted cysts exhibit slightly greater stiffness than wild-type (WT) cysts of the same size. This unexpected finding can be attributed to superelasticity and excess interfacial area of the tissue. E-cadherin-deficient cysts are more strained than WT-cysts as they display highly compressed cells (Fig. 5) (*α* = *h*/*w* − 1), inevitably leading to a larger apparent area compressibility modulus since less areal strain can be mitigated by the superelasticity effect (Fig. 5E). The cells not only recruit additional area from shrinking the thickness of the shell under volume conservation but also from excess interfacial area (Fig. 9B/C). Particularly, the ZO-1 staining accentuates the apical zigzag pattern in WT-cysts indicating a surplus of interfacial area, a feature that is entirely missing in E-cadherin KO cysts (Fig. 8). The surplus area *A*_ex_ inevitably leads to an increasing apparent compressibility modulus as 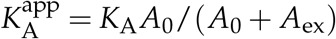. Excess interface area is consequently sacrificed by the tissue to mitigate the decreased stability of adherens junctions (Fig. 9B/C).

In conclusion, our findings reveal that the short-term response to external deformation in curved epithelia is influenced by a complex interplay involving the actomyosin cytoskeleton, cell-cell contacts, and excess interfacial area. Notably, cysts and hemicysts exhibit distinct cell and tissue morphologies, impacting their response to deformation. The observed differences in stiffness and energy dissipation primarily originate from the apical-basal polarity of the cortex in cells constituting the curved tissue. The viscoelastic properties of the actomyosin cortex, coupled with the superelastic behavior of the tissue, play a crucial role in efficiently absorbing external stress. Specifically, we find that the basolateral membrane is significantly stiffer and less fluid compared to the apical membrane facing the lumen, largely due to the greater abundance of excess area apically. Simultaneously, the compliance of cell-cell connections, associated with the cytoskeleton, endows the tissue with Young’s moduli that effectively counteract deformation over brief time periods (seconds). On longer time scales, cell rearrangements further contribute to tissue fluidization under stress. Overall, the morphological response of epithelial tissue to pressure gradients on short time scales strongly depends on the direction of pressure relative to the tissue’s apical-basal polarity.

## MATERIALS AND METHODS

### Cell Culture

Madin-Darby Canine Kidney cells (strain II, MDCK-II; European Collection of Authenticated Cell Cultures, Salisbury, UK) and MDCK-II Cdh1 KO cells were cultured in MEM (Minimal essential medium with Earl’s salts, 2.2 g/L NaHCO_3_, and 2 µM GlutaMAX, ThermoFisher Scientific, Waltham, Massachusetts, USA) with 10% FCS (Fetal Bovine Serum, BioWest, Nuaillé, France) at 37^°^C and 5% CO_2_ in a humidified incubator. The cells were passaged two to three times per week before reaching confluence using phosphate-buffered saline pH 7.4 (PBS; Biochrom, Berlin, Germany) containing trypsin/EDTA (0.25%/0.02% w/v; BioWest/Biochrom, Berlin, Germany). To develop hemicysts, 150,000 cells were seeded in µ-Dishes (35 mm, ibiTreat # 1.5 polymer coverslip; ibidi, Martinsried, Germany) and grown to reach confluence. About 4 days after confluence, hemicysts form spontaneously. To growth cysts, 50,000 cells were seeded in the same media containing additionally 10% of Matrigel (Corning, NY) in µ-Dishes (35 mm, # 1.5 glass coverslip; ibidi, Martinsried, Germany). These dishes were incubated with laminin (0.02 mg/mL, Sigma-Aldrich, Steinheim, Germany) for 2 hrs at 37^°^C prior to seeding. Media is exchanged to MEM with 10% FCS and HEPES (15 µM, Biochrom, Berlin, Germany) for the measurements to remove the Matrigel. Different inhibitors were applied 15 minutes before measurements, latrunculin A (Lat A; 1mM, Sigma-Aldrich, Steinheim, Germany), blebbistatin (Bleb; 50 µM Sigma-Aldrich, Steinheim, Germany), Rock Activator II (CN03; 1 µg/mL, Cytoskeleton Inc., Denver, CO, USA), Y-27632 (RckI; 30 µM, Darmstadt, Germany), Jasplakinolide (Jspk; 1 µM, Enzo Life Sciences, Lörrach, Germany), Calyculin A (CalyA; 20 nM, Fisher Scientific, Schwerte, Germany), and EDTA (5 mM; Biochrom, Berlin, Germany).

### Immunostaining and Confocal MicQQQroscopy

The plasma membrane was stained with CellMask™ (Invitrogen, Carlsbad, CA, USA) and nuclei with Hoechst 33342 solution (Thermo Scientific, Waltham, MA, USA). Cells were kept in MEM for 15 min at 37 °C, afterwards the samples were washed with PBS once before adding MEM media again. For applying polarity markers, samples were carefully washed twice with phosphate-buffered saline (PBS) and fixed with 1 mL of 4 % paraformaldehyde (PFA) solution, kept for 20 min in this solution at room temperature. After that, PFA was carefully aspirated from the samples, and the cells washed three times with PBS after being permeabilized by incubation with 1.0 mL 0.1 % Triton X-100 dissolved in PBS for 10 min at room temperature. Samples then were rinsed twice with 1.0 mL PBS and subsequently incubated with 1.0 mL of blocking solution (2 % (*w/v*) bovine serum albumin (BSA) and 0.1 % Tween 20 (*v*/*v*)). The samples were subsequently incubated overnight with primary antibodies (mouse anti-Podocalyxin (GP135, 1:100, MABS1327, Sigma-Aldrich Chemie GmbH, Taufkirchen, Germany), rabbit anti-Laminin (1:100, L9393, Sigma-Aldrich Chemie GmbH, Taufkirchen, Germany) that were diluted in the blocking solution. Samples were rinsed three times with PBS and then exchanged with the secondary antibodies mixture comprising antibody goat anti-mouse IgG (H+L) Alexa Fluor 546 (1:400, 10002502, Fisher Scientific GmbH, Schwerte, Germany) and goat anti-rabbit IgG (H+L) AlexaFluor488 (1:400, 10729174, Fisher Scientific GmbH,Schwerte, Germany) for 2 hrs at room temperature. After incubation the samples were rinsed three times with PBS and nuclei stained with DAPI (1:10, Sigma-Aldrich, Steinheim, Germany) for 15 min. Afterwards the samples were rinsed three times with PBS and kept in this solution for imaging. For the cell junctions markers, the tissue samples were fixed with PFA as described above followed by permeabilization and blocking. Primary antibody rat anti-UVOMORULIN/E-Cadherin, CLONE DECMA-1, monoclonal (1:500, U3254, Sigma-Aldrich Chemie GmbH, Taufkirchen, Germany) was applied overnight, followed by rinsing the samples three times with PBS, and incubating them with the secondary antibody goat anti-rat IgG (H+L) AlexaFluor 546 (1:400, 10729174, Fisher Scientific GmbH, Schwerte, Germany) in combination with the conjugated antibody mouse anti-ZO1 AlexaFluor 488, monoclonal (10777035, Thermo Fisher Scientific, Waltham, MA, USA) kept for 2 hrs at room temperature and then rinsed three times with PBS. Finally nuclei were stained by adding DAPI (1:10, Sigma-Aldrich, Steinheim, Germany) for 15 min and rinsing the samples three times with PBS, keeping them in solution for imaging.

Imaging was performed with a Confocal Laser Scanning Microscopy (CLSM) (FluoView1200; Olympus, Tokyo, Japan) mounted on an Olympus IX83 Microscope equipped with a 63*×* oil immersion objective (NA= 1.25), acquiring z-stacks with a step size of 1 µm.

### Scanning ion conductance micropscopy

The surface topography of cysts and hemicysts was imaged with SICM using a custom-build setup.^50^ Briefly, fixed cysts and hemicysts samples were imaged in PBS at room temperature in hopping mode using nanopipettes with a typical inner opening radius of 100 nm. The pixel resolution was chosen between 150 and 500 nm and the *IZ*-curve varied between 1 and 3 µm, depending on the imaging area. Images were corrected for sample tilt and offset.

### Atomic Force Microscopy

Force spectroscopy measurements were carried out with a JPK Cellhesion 200 (AFM; Bruker Nano, Berlin, Germany) and a NanoWizard 4 XP (AFM; Bruker Nano, Berlin, Germany) mounted onto an Olympus IX83 microscope. Cantilever CP-FM-BSG-C (NanoAndMore GmbH, Paris, France) with a probe diameter of 20 µm and a spring constant of 0.5 − 2 N/m were used to deform cysts and hemicysts. Force curves were acquired at a speed of 1 µm/s, with a yield force of 20 nN, and a contact time of 5*s* in constant height mode with a sample rate of 4096 Hz. For single cell measurements, MLCT (Bruker AFM Probes, Camarillo, USA) cantilevers with conventional tips were utilized. Yield force was set to 1 nN reached at an approach velocity of 1 µm/s. Force curves were evaluated with a self-written Python script to correct for the baseline, to determine the contact point (Fig. SI7) and to fit the viscoelastic model to the data.

### Computation of indentation-relaxation curves

Three conditions apply to an indented (hemi)cyst that permit computing force-indentation curves 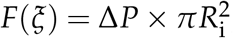, volume conservation and force balance at the top and bottom of the (hemi)cyst, respectively. Essentially, the shape of the deformed epithelia is adopting a minimal area surface as this shape minimizes the free energy (eq. (1)).

### Generic shape function

The dimensionless force 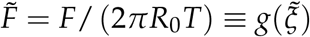 depends only on the shape of the deformed cyst, with the dimensionless indentation depth 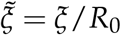. Consequently, for cysts 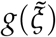 depends on 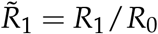 and 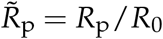, while for hemicysts the function depends on the contact angle *ϕ* and 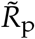. The shape represented by 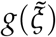 is approximated with a polynomial to 4^th^ order providing an analytical fitting function 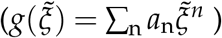. The same is done for the area expansion 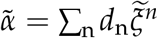 of the cyst during indentation. The generic shape function 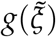 as well as the corresponding area expansion 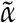 can be computed for a given indenter radius *R*_p_ and angle *ϕ*.

### Viscoelastic solution

Viscoelasticity is considered by assuming that the 2D elastic modulus obeys a power law 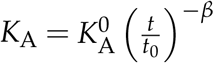 with 0 ≤ *β* ≤ 1 and *t*_0_ = 1 s (set arbitrarily).^13, 51^ Application of the elastic-viscoelastic-correspondence principle leads to the following expression for the overall tension:^13, 19^

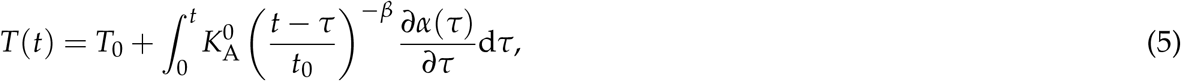

with *α* = Δ*A*/*A*_0_. The hereditary integral (eq. (5)) can be solved analytically by Laplace Transformation. The force response of cysts to indentation or compression (approach) at constant velocity 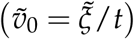 gives:

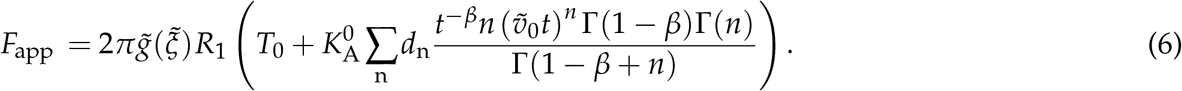

The force response upon relaxation starting at *t* = *t*_m_ with zero velocity is:

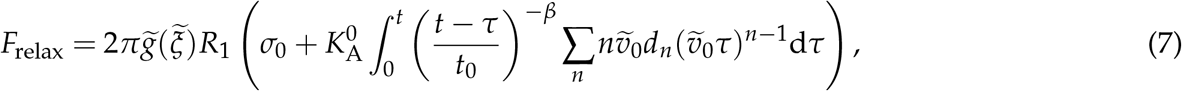

with the Gamma function 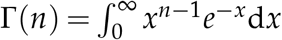. The corresponding generic shape function, 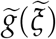, needs to be chosen according to the geometry of cyst and indenter. Experimental indentation-relaxation curves were subject to fitting a piecewise function *f* (*t* ≤ *t*_m_) = *f*_app_ (*t*) and *f* (*t* > *t*_m_) = *f*_relax_(*t*). In the following, the term area compressibility modulus or stiffness refers to the scaling factor of the area compressibility modulus 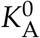, while the power law exponent *β* designates the fluidity of the hemicyst/cyst.

### Superviscoelastic solution

Laplace pressure causes the thickness of the curved cell monolayer to decrease, thereby releasing lateral tension and reducing the overall tension in the shell (Fig. SI5).^27^ The apical/basal tension balancing the pressure gradient across the cell monolayer is partially compensated by thinning of the cell layer as depicted in Fig. SI3-6. This gives rise to an expression for the pre-stress 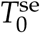, which depends non-linearly on the area dilatation *α*_0_:

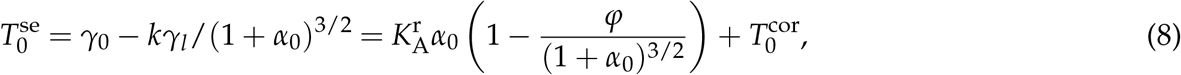

with 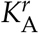, the area compressibility modulus of the curved epithelia, 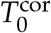 the cortical tension that does not depend on the areal strain and 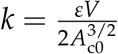, a constant capturing the geometry of the cells. *V* is the volume of the cell, *A*_c0_ is the apical/basal area prior to strain, and *ε* a geometric parameter (see SI chapter 1.7). *γ*_*l*_ = *ϕ*(*γ*_0_ − *γ*_ad_) is the lateral tension of the cell comprising the lateral cortical tension *γ*_*l*_ reduced by adhesion energy per unit area *γ*_ad_ due to formation of cell-cell contacts.^27^

### Apparent area compressibility modulus

External forces lead to additional area dilatation of already pre-stressed shells. The overall tissue tension including superelasticity can then be written as a linear combination of the pre-stress above and the areal strain due to indentation 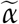 times the apparent area compressibility modulus 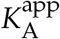:

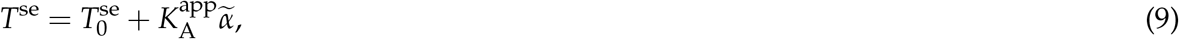

with 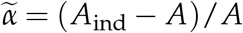 the areal strain during indentation with the new increase surface area after deformation and the unperturbed, but pre-stressed area *A*. Combining these equations gives:

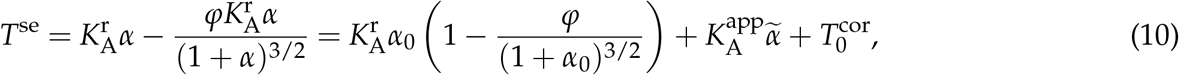

with *α* = (*A*_ind_ − *A*_0_)/*A*_0_, the areal strain during indentation with the new surface area *A*_ind_ and the unperturbed and unstressed surface area *A*_0_. Note that *α*_0_ is substantially larger than 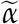 as *A* ≫ *A*_0_. Consequently, the apparent area compressibility modulus measured in indentation experiments reads:

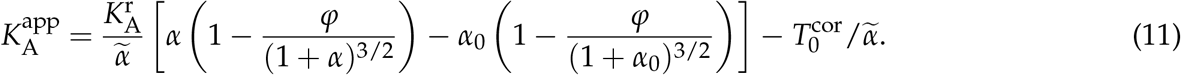

As a consequence, low pre-stress, synonymous with a small initial strain *α*_0_, leads to lower apparent stiffness.

### Statistical Analyses

For statistical analysis the data sets were compared using the non-parametric Mann-Whitney-Wilcoxon rank sum test (ns: *p* value > 0.05, * denotes *p* < 0.01, ** represents *p* < 0.001, and **** indicates *p* < 0.00001). Experiments were performed over different days with different cell samples.

## Supplementary Information

### 1 Theory

The alteration in the shape of a shell structure occurs through two primary mechanisms: in-plane stretching and shear or out-of-plane bending. The significance of these deformation modes and the extent to which they are permitted dictate the development of various theoretical frameworks, each tailored for specific scenarios.

The force response of curved epithelia to indentation is described below. The following assumption were made:

- Bending is neglected. This is justified since large curvatures are avoided by using spherical indenters and the shell is thinner than the radius of the domes of cysts.
- The thickness of the cyst is considered to be infinitely small. This approximation goes hand in hand with ignoring bending contributions. This assumption is partly given up when implementing superelasticity.

#### 1.1 Young-Laplace equation of spherical vessel

To estimate the ramifications of our assumption (thin shells) we will first revisit the derivation of Young-Laplace’s equation for a spherical cyst of thickness *h* (height of the cells).

**Figure SI1:**
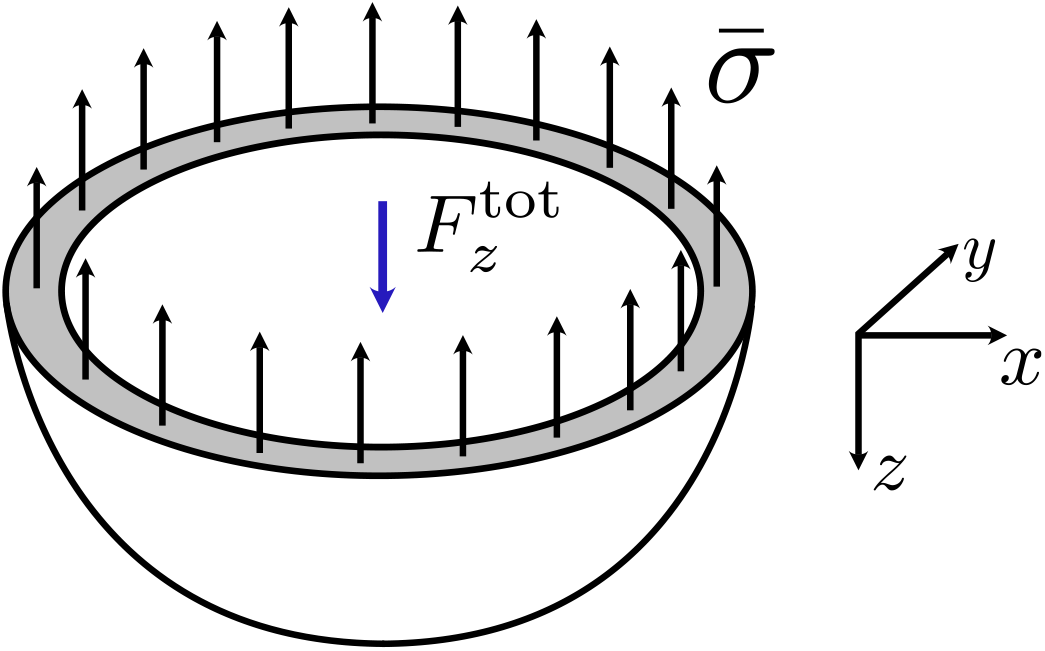
Force balance of a cyst with thickness *h*, i.e., forces are compensated by internal stresses 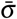.

We set the center of the cysts, with the inner radius of the deformed geometry denoted by *r* and the outer radius by *R*. The force per unit area acting on the inner surface of the sphere can be cast in spherical coordinates (*θ*, *ϕ, r*):

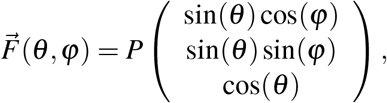

with *θ* ∈ [0, *π*] and *ϕ* ∈ [0, 2*π*] and *P*, the pressure acting on the cyst. The obtain the overall forces in *x, y*, and *z* direction, respectively, they need to be integrated over the surface of the cyst:

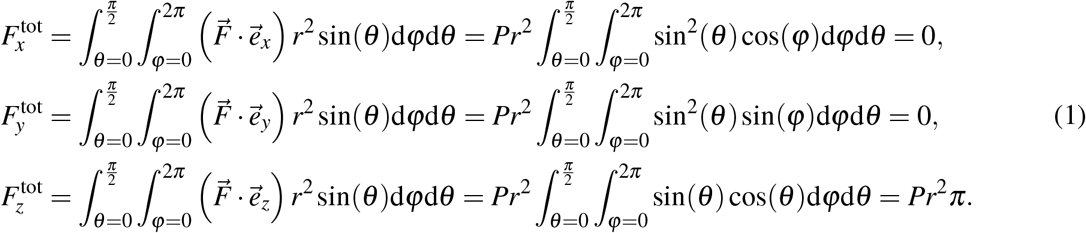

We only need to integrate over the upper half of the cyst, hence we have 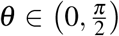. Forces in *x* and *y* direction vanish. Assuming homogeneous stress, we approximate the wall stress *σ* by its mean value 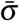. The tangential components are identical in any direction since the geometry and the acting forces are symmetric and therefore shear stress vanishes. Taken together, the mean wall stress at the cross-section acts solely in *z*-direction, thus we have 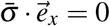 and 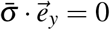. The three components of the total wall stress vector 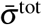 are:

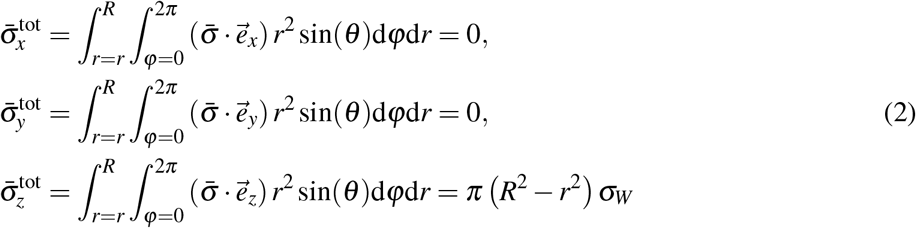

with 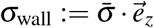.

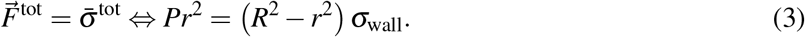

This gives:

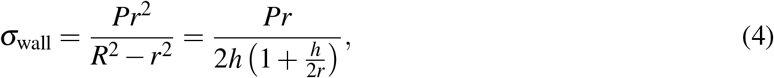

with *h* the thickness of the shell. Assuming *h* ≪ *r* we obtain:

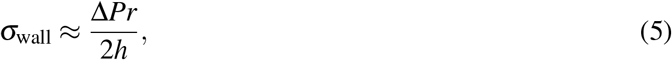

with *T* = *σ*_wall_*h* and *r* = *R* we arrive at the well-known Young-Laplace equation for a sphere:

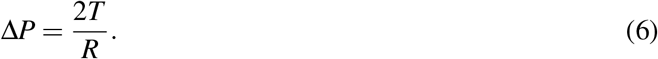

#### 1.2 Elastic description of a planar cell sheet under biaxial strain

Here, we initiate our analysis with the aforementioned thin film approximation, under the assumption that all forces involved are conservative (later we will employ the elastic-viscoelastic correspondence principle to integrate dissipation). This simplification leads us to describe the uniform tension according to a 2D Hookean law:

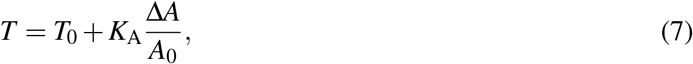

with *T*, the tension and *K*_A_ the area compressibility modulus. Δ*A* = *A*− *A*_0_ is the area change and *A*_0_ the unperturbed area. If *T*_0_ originates from pre-stress *T*_0_ = *K*_A_ (*A*_0_ − *A*_i_)/*A*_i_ = *K*_A_*α*_0_, with *A*_*i*_, the initial area at zero Laplace pressure, we have to write:

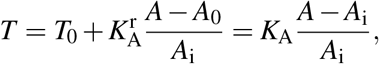

with 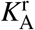, the real area compressibility. This means that

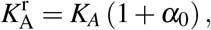

with *α*_0_ = (*A*_0_ −*A*_i_)/*A*_i_. This makes a relevant contribution only if the initial area dilation is substantial.

#### 1.3 Deformation of spherical cysts with a spherical indenter

##### 1.3.1 Shape of the cyst during indentation

The cyst is assumed to be formed by a sufficiently thin sheet, which permits us to neglect bending and shearing contributions to the free energy functional 𝔉. Minimizing the free energy

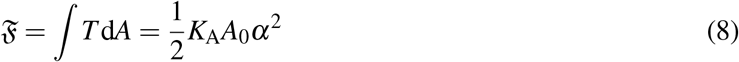

under constant volume assumption is therefore identical to finding the minimal surface of the compressed cyst. *A*_0_ is the area of the unstressed cyst, *T* the tissue tension, *K*_A_ is the area compressibility modulus and *α* the areal strain 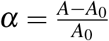. Solutions are contours of constant curvature. Assuming rotational symmetry around the *z*-axis, minimizing the free energy is identical to minimization of the area under constant volume. This provides the following Euler-Lagrange equation:

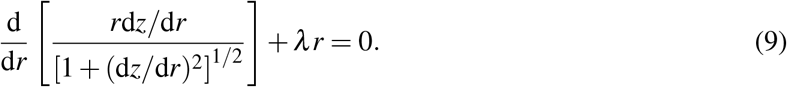

The reciprocal principal radii of curvature are

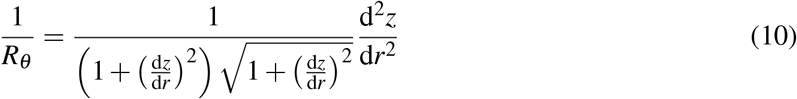

and

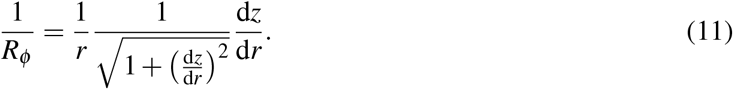

*R*_*θ*_ and *R*_*ϕ*_ denote the meridional and circumferential radii of curvature at any point in the membrane. Comparing equation (11) with equations (12) and (13) permits us to conclude that *λ* represents the constant curvature of the cyst:

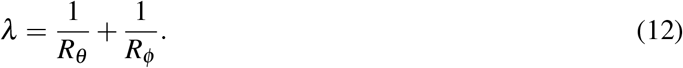

Compared with mechanical equilibrium for normal forces in the absence of shear, *λ* is identical to the pressure difference across each surface element divided by the isotropic tension, giving rise to the Young-Laplace equation:

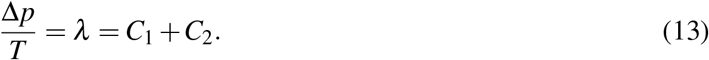

with the principle curvatures 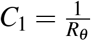 and 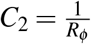 at each point of the freestanding parts of the (hemi)cyst (see Fig. SI2). The principle curvatures can be expressed in terms *u*(*r*) = sin *γ* and *r*:

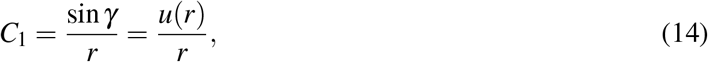

and

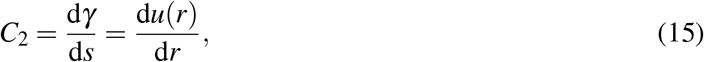

where *s* is the curvilinear distance along the surface meridian and *γ* the angle with the surface normal. Note that 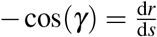 as obtained from differential geometry. We can now rewrite the Young–Laplace equation as the total *constant* curvature:

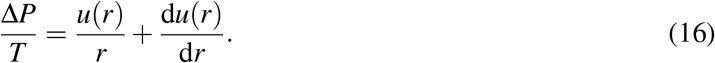

This relation can easily be integrated:

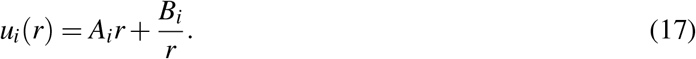

with *i* = 1, 2, 3 referring to the corresponding regions of the free contour (*s*_1_ → *s*_2_(*i* = 1), *s*_2_ → *s*_3_(*i* = 2), *s*_3_ → *s*_4_(*i* = 3)). For each of the regions appropriate boundary conditions have to be fulfilled. *A*_1_ and *B*_1_ correspond to the unbound region *i* = 1 ranging from *s*_1_ → *s*_2_. The following boundary conditions hold:

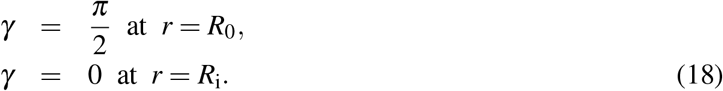

where *R*_*i*_ is the contact radius of the cyst formed with the flat substrate at the bottom and *R*_0_ the equatorial radius of the deformed cyst (see Fig SI2). From Eqs (17, 18) we obtain,

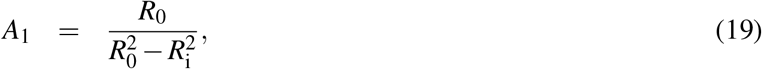

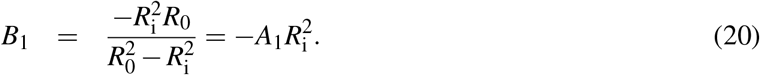

In region *i* = 2 (*s*_2_ → *s*_3_) the free contour obeys the boundary conditions:

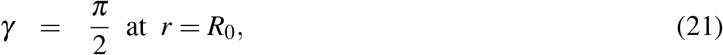

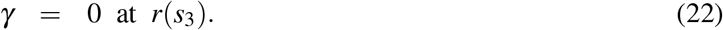

**Figure SI2:**
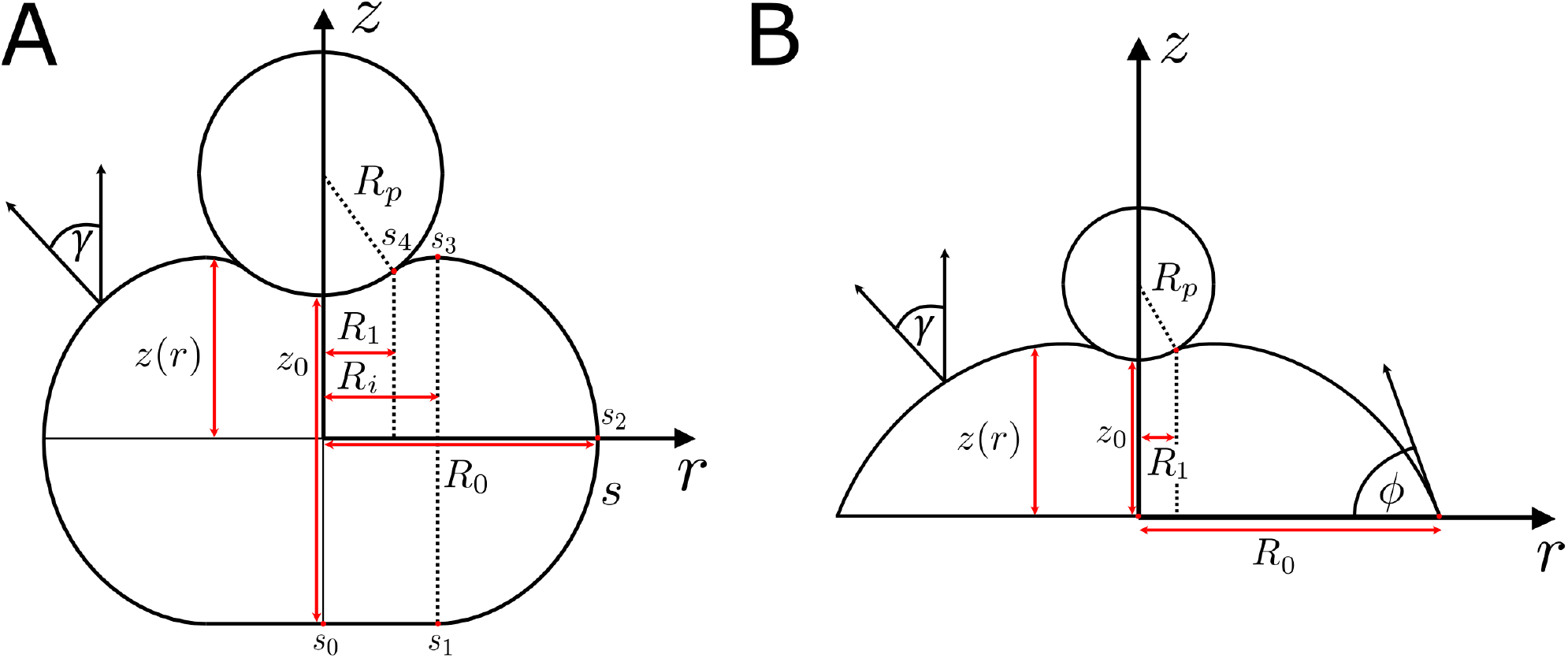
Parametrization of a cyst (A) and hemicyst (B) indented with a spherical indenter of radius *R*_*p*_.

Therefore,

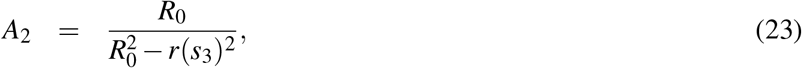

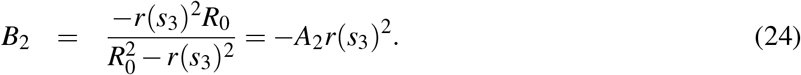

Since the contour is continuous at *R*_0_, *r*(*s*_3_) = *R*_i_ holds and therefore also *A*_1_ = *A*_2_ and *B*_1_ = *B*_2_, i.e. the free contour from *s*_1_ → *s*_2_ and *s*_2_ → *s*_3_ are mirror-inverted. *A*_3_ and *B*_3_ for region *i* = 3 (*s*_3_ → *s*_4_) that reaches up to the contact with the indenter at *R*_1_ are obtained from the following boundary conditions:

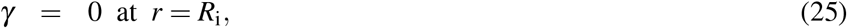

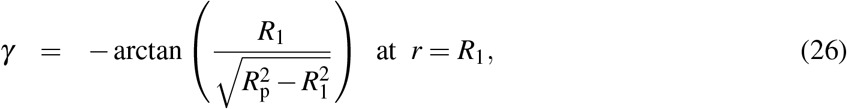

leading to

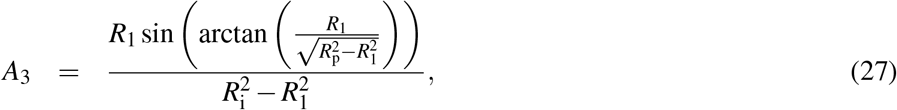

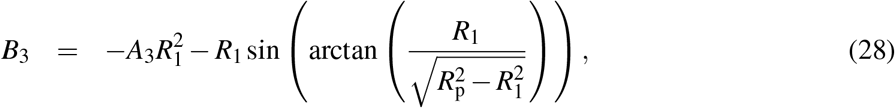

where *R*_p_ is the radius of the indenter. Once the radii *R*_0_, *R*_i_, and *R*_1_ are found, the free contour corresponding to the regions (*s*_1_ → *s*_2_ using *u*_1_(*r*), *s*_2_ → *s*_3_ using *u*_2_(*r*) = *u*_1_(*r*), and *s*_3_ → *s*_4_ using *u*_3_(*r*)) can be readily obtained from integrating:

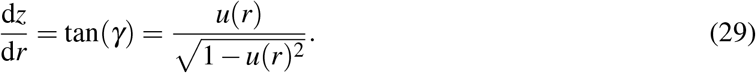

The remaining contour is defined by the boundaries, a flat substrate at the bottom and the conical indenter from the top.

The task is now to find expressions for *R*_0_, *R*_1_, and *R*_i_ depending on the distance between the tip of the indenter and the flat base plate at the bottom *z*_0_. Three conditions apply to an indented cyst that permit computing force-indentation curves (*F*(*ξ*)). The following section describes how to find a set of parameters *R*_0_, *R*_1_, and *R*_i_ at a given force.

##### 1.3.2 Force balance and volume constraint

The volume of the cyst prior to indentation is denoted as *V*_v_ and the volume of the indented one *V*_ind_. The condition of volume conservation reads:

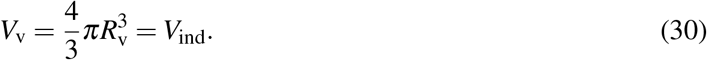

The indented cyst is a solid of revolution, which facilitates the integration to obtain the volume *V*_ind_:

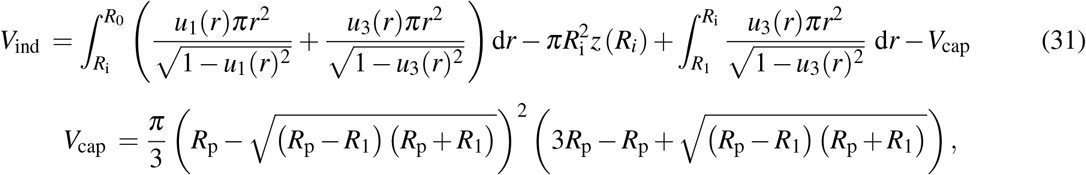

and 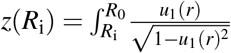.

The key assumption is that the only source of the restoring force to indentation is the in–plane tension 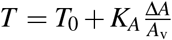 due to area dilatation. Δ*A* = *A*_ind_ − *A*_v_ denotes the difference between the actual area *A*_*ind*_ and the initial area prior to compression *A*_v_. *T*_0_ is the tension of the cyst. The force balance of the top part of the cyst in the *z*-direction has two contributions, one from a blunt indenter with a radius of *R*_1_ and a sphere immersed with conformation contact with the cysts 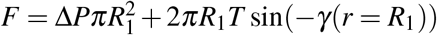:

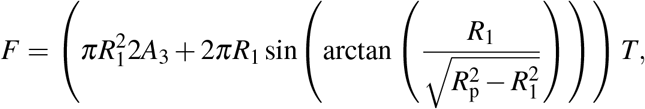

leading to

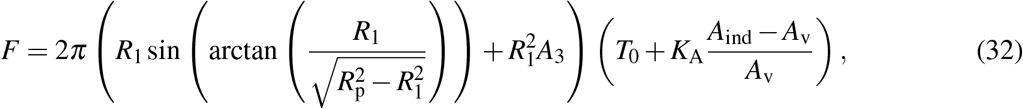

which is the second condition, while force equilibrium at the bottom part is the third condition:

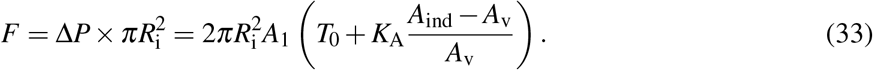

###### Surface area *A*_ind_ of the cyst

The area *A*_v_ prior to indentation is 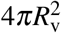, while the actual area *A*_ind_ can be divided into two surfaces of revolution, the top 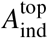 and bottom part 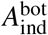 of the cyst according to Fig SI2:

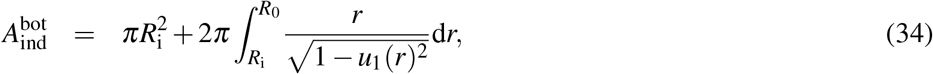

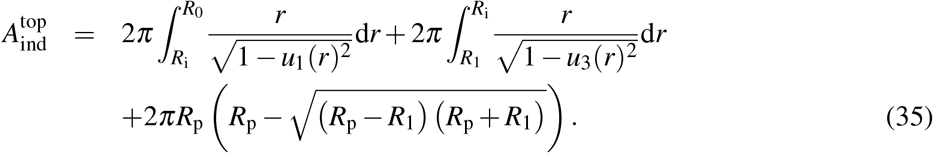

###### Indentation depth

The indentation depth *ξ* in the center at *r* = 0 is readily obtained from:

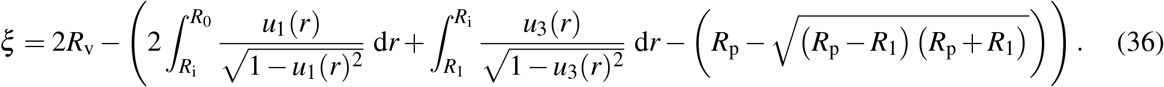

The contour in region *s*_1_ → *s*_3_ corresponds to the first integral, while the contour along the path *s*_3_ → *s*_4_ is represented by the second integral.

##### 1.3.3 Stretching versus bending

When a thin, water-filled capsule under constant volume is deformed by a spherical indenter, the resultant restoring forces arise from in-plane stresses, namely the stretching of the capsule’s exterior, shear stress, and out-of-plane bending. In order to simplify the analysis, we first determine whether the Hamiltonian is dominated by bending or stretching of the shell. Provided that the shell’s thickness *h* is adequately small relative to its radius *R*, we can express the ratio of the stretching energy per unit area to the pure bending energy per unit area as follows:[1, 2]

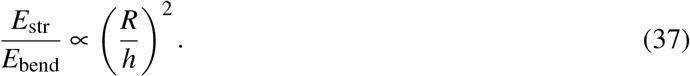

This ratio tells us whether the response of thin walled capsules to deformation is governed by stretching or bending. Cysts in our case possess an approximate ratio 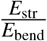 of 10-100 in favor of a stretching-dominated regime. As spherical capsules inevitably can not be deformed without stretching, area dilatation dominates the response to indentation. A more thorough analysis of Couturier et al.[3] shows for small indentation depths (*ξ*) and small contact regions that the restoring force to indentation with a spherical indenter is independent of the indenter size and leads to a linear relationship between force and indentation depth:

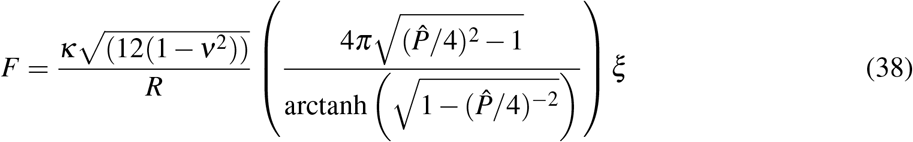

with 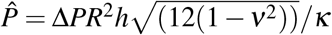, in which *κ* = *E*_Y_*h*^3^/(12(1−*ν*^2^)) denotes the bending modulus of the shell with *E*_Y_ the Young’s modulus and *ν* the Poisson ratio. The pressure difference between lumen and the outside is related to the tension *T* by Laplace’s law (equation (**??**)). In the limit of an unpressurized shell, the linear relationship between indentation and force of Reissner’s classical shell theory for shallow spheres is recovered 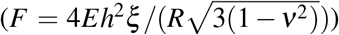 suggesting a linear relationship between force and indentation.[4, 5, 2] Equation (38) predicts a decrease of the sample stiffness *k*_s_ = d*F*/d*ξ* with the radius of the curved epithelia. Since the pre-stress governing the response to low strain increases as the radius of the curved epithelia decreases, we cannot entirely rule out the contribution of bending to the low strain response. Even if the pressurized spherical shell behaves like a linear spring under small compression due to bending, this effect will be largely overshadowed by the pre-stress term (see Fig. SI3). To demonstrate this further we computed the various contributions to the Hamiltonian as a function of areal strain using a minimal area surface where *E*_str_ is minimal. For the contribution of bending this is an upper approximation. Fig. SI3 shows that highly curved epithelia and low pre stress show bending dominated response at low areal strain. In our experiments, this case is largely avoided as reasoned below. Shear is also excluded as a major contribution to the stiffness of the cysts as the cysts is globally pressurized and not experiencing substantial shear stress compared to the in-plane expansion. In the following, we will compare the different contributions to the Hamiltonian.

**Figure SI3:**
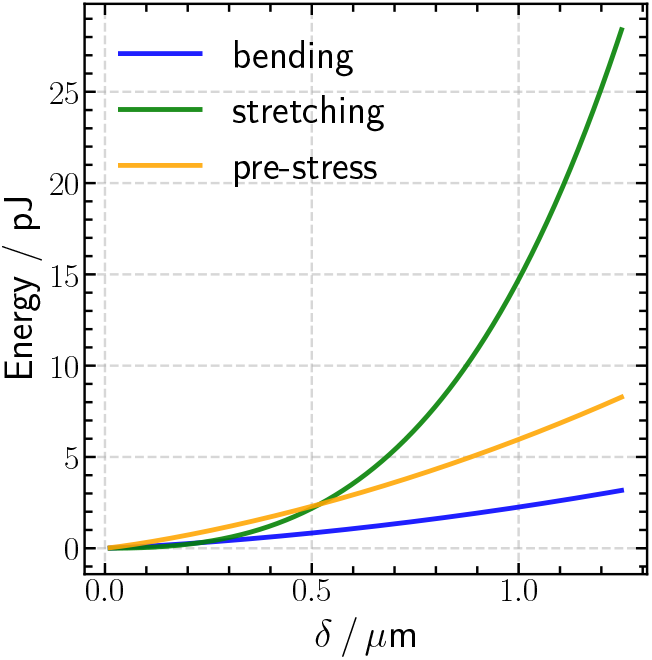
Comparison of the different conceivable energy contributions to deformation of cysts subject to external force exerted by a spherical indenter. Thickness of the shell: *h* = 4.5 *µ*m, Poisson’s ratio: *ν* = 0.33, radius of the cyst: *R*_0_ = 30 *µ*m. Area compressibility modulus: *K*_A_ = 0.15 N/m. Pre-stress: *T*_0_ = 4 mN/m.

Assuming that the cells organized in curved epithelia obeys Hooke’s law and are isotropic, i.e., characterized by two material constants like Young’s modulus *E* and Poisson’s ratio *ν*, for thin shells the strain tensor is only related to the effective membrane stress tensor and the bending tensor only to the effective moment tensor. Proportionality constants are the area compressibility modulus *K*_A_ and the bending modulus *κ* = *K*_A_*h*/(12(1 − *ν*^2^)) In the following we will show when we can neglect bending. The change in stretching energy can be computed from

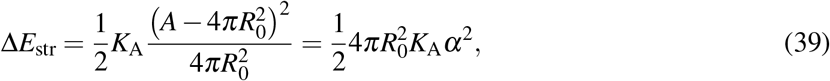

with the stretched area *A* upon indentation:

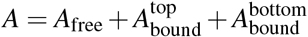

compiling the free area:

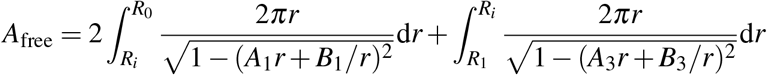

and the bound ones to the probe (top) and the substrate (bottom), respectively:

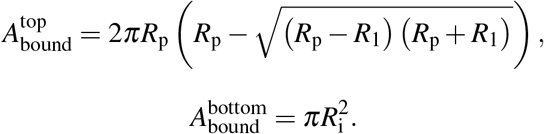

The energy of bending is obtained from

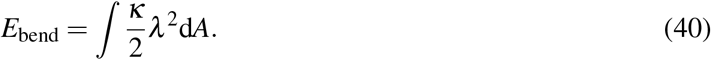

The observed behavior in cysts and hemicysts primarily reflects the mechanical properties of individual cells, specifically their apical-basal polarity. In cysts, the basal side of cells facing outward is less compliant than the apical side, partly due to the excess surface area of the apical side with the extrinsic curvature *λ* = *C*_1_ + *C*_2_ and the bending modulus 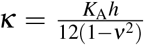, *ν* is the Poisson number. Now, we compute the bending energy for the same minimal surface obtained from solving Young-Laplace’s equation. This is a convenient approximation as *λ* = const and any true bending energy from solving the Euler Lagrange equation originating from the full Hamiltonian will be smaller. As *λ* = 2*A*_1_ we arrive at the change of free bending energy:

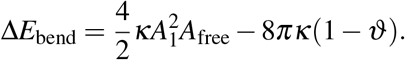

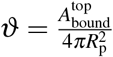 is the fraction of the probe covered with the cells during indentation.

The energy contribution due to pre-stress is simply obtained from:

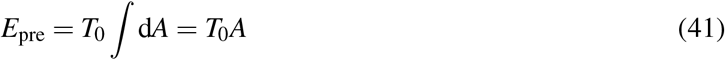

As the base radius of hemicysts is substantially larger than the radius of the cysts, we can exclude bending in the energy functional describing deformation of domes.

#### 1.4 Deformation of hemicysts/domes with a spherical indenter

##### 1.4.1 Shape of the domes during indentation

The indented hemicyst is parameterized as shown in Fig. SI2. Essentially, the non-deformed shape is a spherical cap with contact angle *ϕ*_0_ and base radius *R*_*i*_ The Young-Laplace equation is solved for the following boundary conditions:

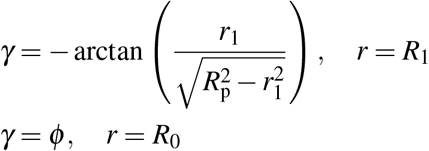

The shape function *u*(*r*) can be written as:

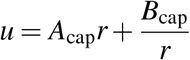

with

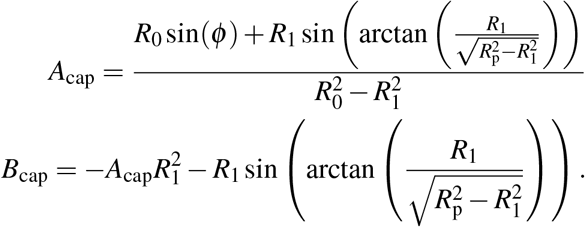

##### 1.4.2 Force balance and volume constraint

The volume can be computed from the shape as a function of indentation depth:

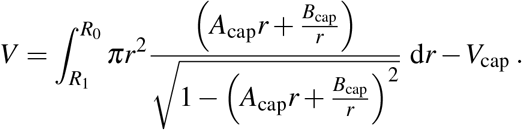

*V*_cap_ is the volume of the spherical cap/dome immersed in the cell upon indentation:

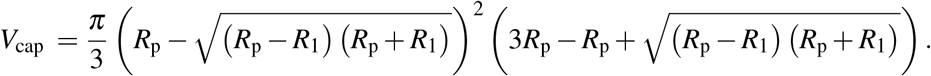

*V* is set equal to the volume of the cell prior to indentation (shape of a spherical cap/dome):

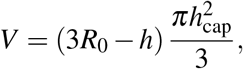

with *h*_cap_, the height of the non-deformed cap/dome.

###### Surface area *A*_ind_ of the dome

The actual area *A* of the indented cell can be readily obtained by solving the integral (surface of revolution):

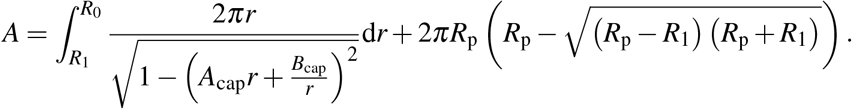

###### Indentation depth

Again, the two non-linear equations are solved numerically for [*R*_1_, *ϕ*]. The *z*-position of the AFM-tip at *r* = *R*_1_ is calculated from:

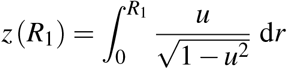

that allows us to determine the indentation depth *ξ* :

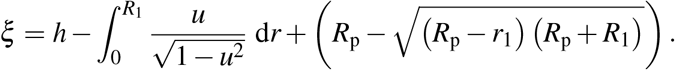

As above, force equilibrium at the base provides the force as a function of *ϕ* and *r*_1_ :

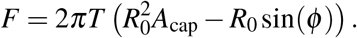

Note that *R*_0_ = *R*_hc_ the base radius of the hemicysts, which does not change.

#### 1.5 Generic shape functions

First, we write everything in dimensionless quantities. The force 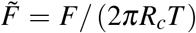 depends only on a function 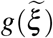 (with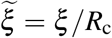) that represents the generic shape of the deformed hemicyst or cyst. *R*_c_ is the initial radius of the cysts or the base radius if the hemicyst (*R*_hc_), respectively. For instance, for the cysts we obtain:

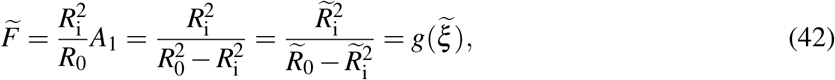

with 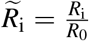 and 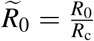 is computed once and can later be scaled for any radius of the cyst and the indenter shape, respectively. 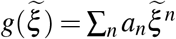 is approximated with a polynomial that provides an analytical fitting function. The same is done for the area expansion 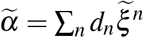 of the cyst during indentation. For the hemicysts we obtain:

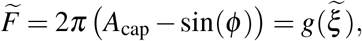

as *R*_0_ = *R*_hc_ for hemicysts. The generic shape functions for cysts and hemicyst, respectively, 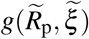 as well as the corresponding area expansion 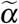 can be computed once for a given set of parameters *ϕ* and *R*_p_, respectively. The numerically computed shape functions can then be approximated by a polynomial to speed up the computation.

#### 1.6 Viscoelastic solution

Viscoelasticity of cyst is entering through the area compressibility modulus of the shell. We assume that this 2D elastic modulus obeys a power law 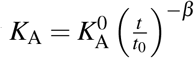 with 0 ≤ *β* ≤ 1 and *t*_0_ = 1s (set arbitrarily).

Application of the elastic-viscoelastic-correspondence principle leads to the following expression for the overall tension:

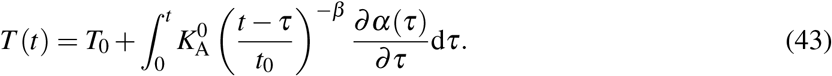

The hereditary integral can be solved analytically by Laplace transform if the corresponding area integrals are also approximated with a polynomial 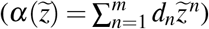.[6] The force response of cysts to indentation or compression (approach) at constant velocity 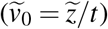 gives:

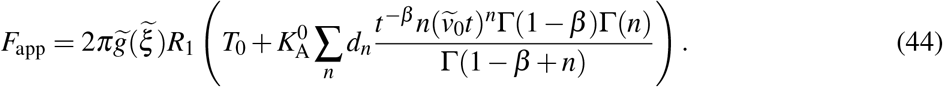

The force response upon retraction at *t* = *t*_m_ with the identical velocity is:

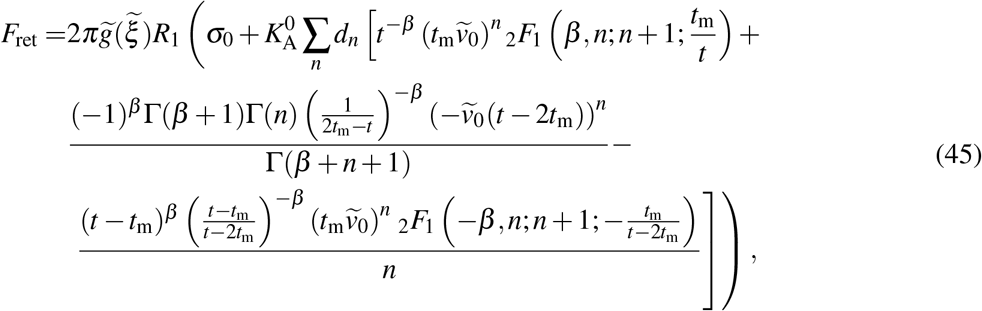

and for relaxation setting in at *t* = *t*_m_:

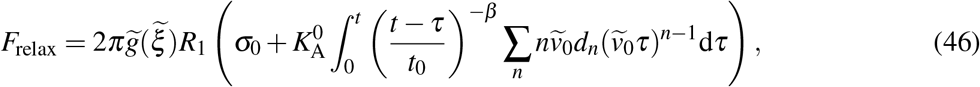

with the Gamma function 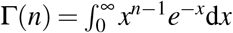. The corresponding generic shape function, 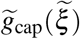, needs to be chosen according to the geometry of cyst and indenter. Usually, polynomials to the order of *m* = 4 are sufficient to describe the functions 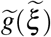 and 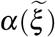 with good accuracy. Experimental force - time curves were subject to fitting a piece-wise function *f* (*t*≤ *t*_m_) = *F*_app_(*t*) and *f* (*t* > *t*_m_) = *F*_ret_(*t*). In the following, the term cyst stiffness refers to the scaling factor of the area compressibility modulus 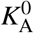, while the power law exponent *β* designates the fluidity of the cyst.

#### 1.7 Superviscoelastic solution

The following treatment is inspired by the work of Trepat and coworkers.[7] First we start again with the free energy of the cyst but this time consider the special shell-geometry consisting of a monolayer of cells that maintain their volume.

**Figure SI4:**
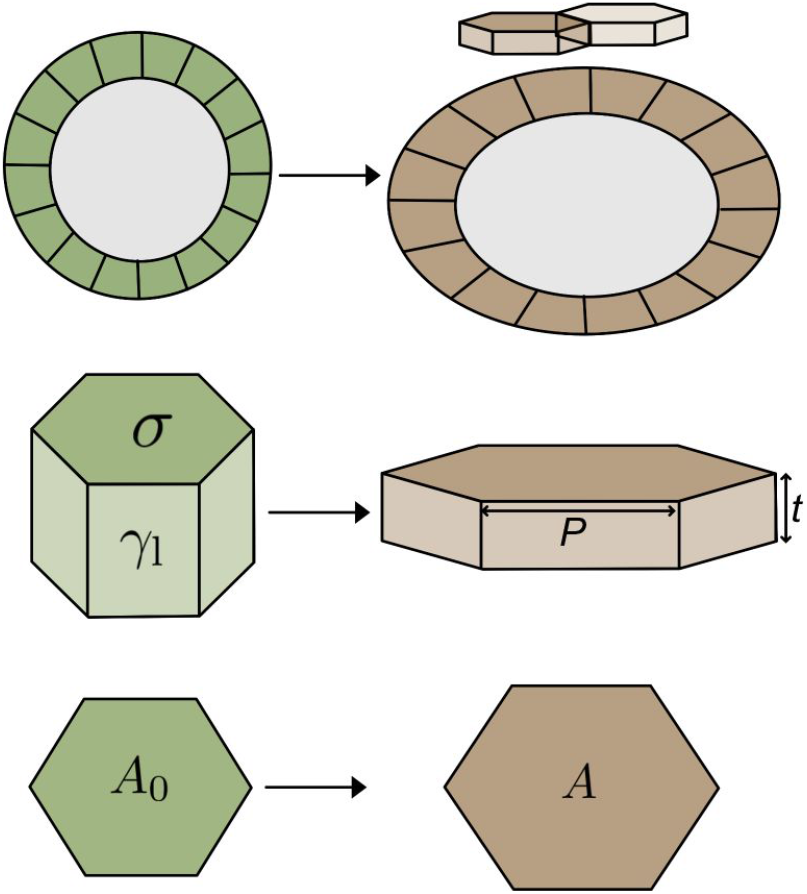
Deformation of a cyst consisting of regular hexagonal cells undergoing uniform areal stretching. We employ the idealized case where all cells in the tissue are assumed to be identical and undergoing the same deformation.

Let *P*_1_ and *P*_2_ be the pressures in the lumen and the outside environment, respectively. Then the work necessary to bring about the change in volume is *W*_vol_ = − ∫ (*P*_1_− *P*_2_) d*V*. The total work or free energy 𝔉 in displacing the surface is obtained by adding to this the work connected with the change in area of the surface. This part of the work *δW*_t_ is proportional to the change *δ A* in area of the surface, and is Σ*δ A*, where Σ is the tissue tension. We obtain:

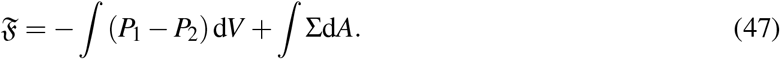

Minimizing the free energy with respect to displacement *ξ* requires:^1^

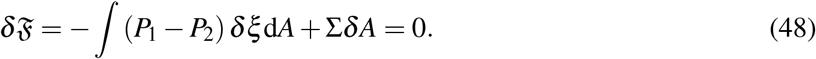

The condition for thermodynamical equilibrium is, of course, that *δ*𝔉 = 0. Next, let *R*_1_ and *R*_2_ be the principal radii of curvature at a given point of the surface. We set *R*_1_ and *R*_2_ as positive if they are drawn into medium 1 - the lumen of the cyst. Then the elements of length (the arc differentials) d*s*_1_ and d*s*_2_ on the surface in its principal curvature sections are increased to (*R*_1_ + *δξ*) d*s*_1_/*R*_1_ and (*R*_2_ + *δξ*) d*s*_2_/*R*_2_ when the angles d*s*_1_/*R*_1_ and d*s*_2_/*R*_2_ remain constant, i.e., an expansion normal to the surface (d*s*_1_ is the arc length of a circle with radius *R*_1_, and correspondingly for d*s*_2_). Hence the surface element d*A* = d*s*_1_d*s*_2_ becomes, after displacement,

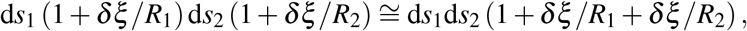

i.e., it changes by *δξ* d*A* (1/*R*_1_ + 1/*R*_2_).^2^ Hence we see that the total change in area of the surface of separation is

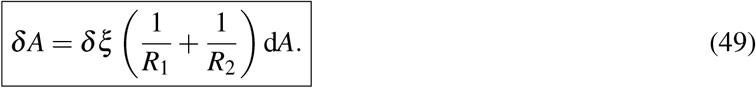

**Figure SI5:**
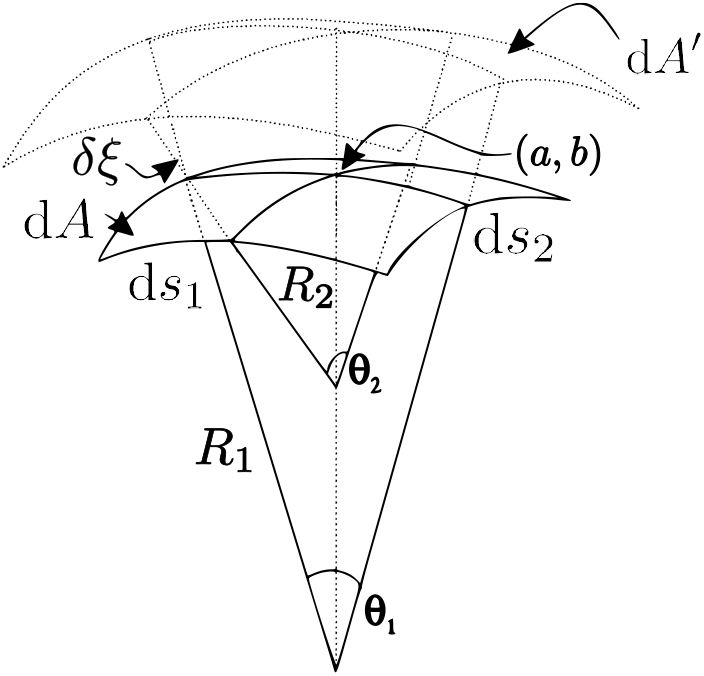
Parametrization of a surface element subject to deformation *δξ*.

Substituting these expressions into the equation for the free energy and equating to zero, we obtain the equilibrium condition in the form

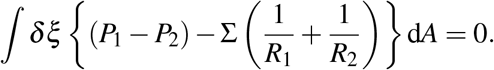

This condition must hold for every infinitesimal displacement of the surface, i.e., for all *δξ*. Hence the expression in braces must be identically equal to zero and Young-Laplace’s equation follows.

Let now revisit the second contribution *W*_t_ to the free energy 𝔉 = *W*_vol_ + *W*_t_. Because of the arrangement of cells within the shell, depicted in Fig. SI4, and their organization in an idealized planar geometry, we can express the total differential of the free energy per unit area as a function of relative area dilatation:

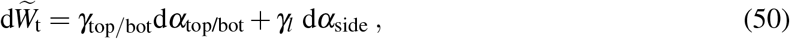

with *α*_top/bot_, the area dilatation of the apical and basal side. *α*_side_ denotes the relative areal change (compression) of the lateral side. *γ*_top/bot_ denotes the apical/basal tension, while *γ*_*l*_ refers to the lateral tension being on the order of *γ*_top/bot_ (see below).[7] For the area dilatation of the top/bottom area and the side we obtain

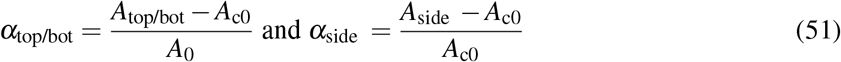

**Figure SI6:**
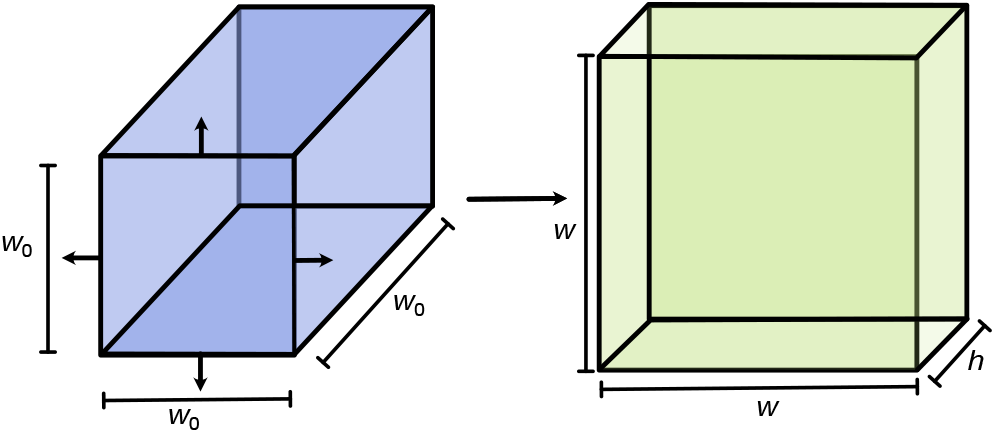
Deformation of a single cell modeled as a cube during stretching of the shell, i.e., undergoing uniform areal stretching under constant volume.

Now we use the volume constraint to obtain *A*_side_ as a function of *A*_top/bot_. A simple derivation of the tissue tension can be accomplished by assuming that cells adopt a cubic morphology prior to stretching.

Assuming volume conservation, we obtain 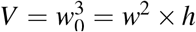, with *w*^2^ = *A*_top/bot_ = *A*_c0_ + Δ*A*, the apical (or basal) surface of the cell. 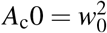 is the initial apical or lateral surface area. This is identical to the surface of the whole cyst *A* divided by the number of cells *n*. The thinned side (thickness *h*) of the cell is then 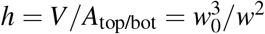. The area of the side is 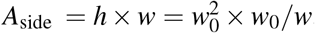, hence shrinks with increasing area dilatation of the top/bottom. Hence we obtain for the area dilatation:

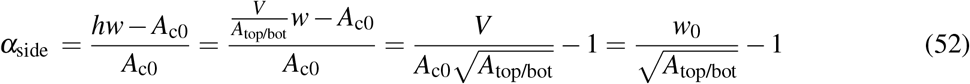

and

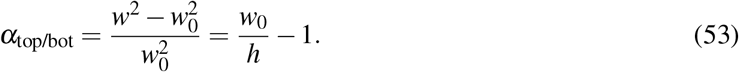

Next we substitute *α*_side_ with *α*_top/bot_ :

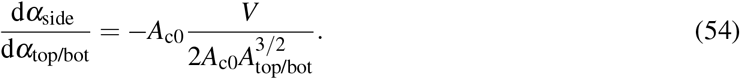

Taken together we obtain for the change in free energy per unit area:

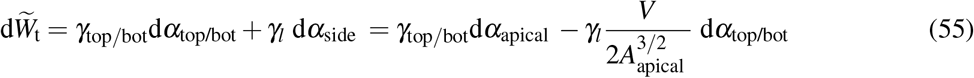

or in terms of *α*_top/bot_

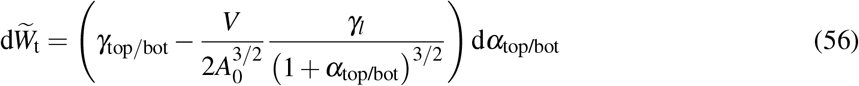

The tissue tension 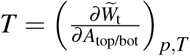 reads generally:

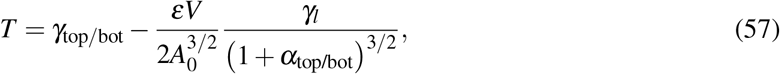

with *ε* = 1 for the cubic cell and for a hexagonal prism we have 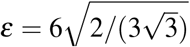.[7] Consequently, the tissue tension *T* is the apical/basal tension *γ*_top/bot_ reduced by the lateral tension *γ*_*l*_ as additional area is provided by thinning of the cell layer under constant volume. The larger the area dilatation, the smaller this effect. We show experimentally that indeed thinning of the cell layer occurs, damping the strain on the cell (Fig. SI6). Assuming that the ratio of *γ*_top/bot_/*γ*_*l*_ is approximately constant, we can write a superelastic tension (*kγ*_*l*_ = *φγ*_top/bot_) as:[7]

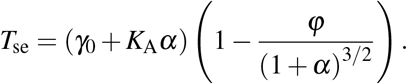

*γ*_0_ is the apical/basal tension prior to indentation. The lateral tension comprises also the adhesion between cells with *γ*_*l*_ = *φ* (*γ*_0_ − *γ*_ad_) with *γ*_ad_, the adhesion energy per unit area. The initial area dilatation *α*_0_ resulting from Laplace pressure is substantial, typically around 50%, whereas the area dilatation coefficient, denoted by *α* = (*A*_n_ − *A*)/*A*, remains quite small, typically on the order of 1%, especially in the context of indentation relaxation experiments. Consequently, the manifestation of superelasticity becomes apparent primarily through an effective pre-stress parameter, denoted as *T*_0_, acting on the pressurized tissue:

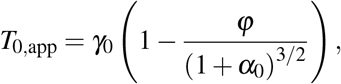

and an apparent area compressibility modulus:

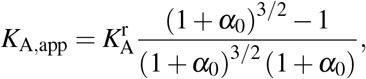

assuming *φ*≈ 1. For the viscoelastic case, we obtain:

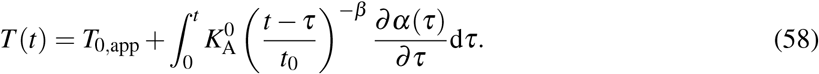

In the following, we will dismiss the index “app” and refer simply to pre-tension *T*_0_.

#### 1.8 Correction factor for large initial areal strain

The following section shows how to correct the area compressibility modulus to fulfill:

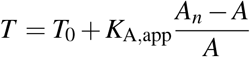

The final tension of the shell is obtained from:

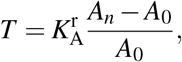

while the initial tension (pre stress) *T*_0_ is:

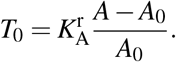

So as we want to compute the final tension as:

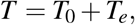

we obtain for *T*_*e*_ :

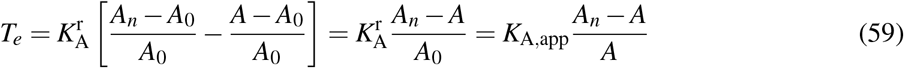

with the measured value

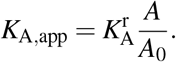

or for the intrinsic area compressibility modulus:

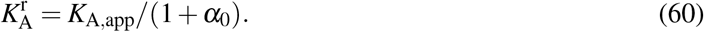

### 2 Automatic detection of the contact point in indentation relaxation data

The algorithm (illustrated in Fig. SI7) enables us to automatically determine the contact point between indenter and epithelia through a series of steps. Firstly, we identify the relevant local regime by setting appropriate thresholds (A), then a knee-elbow algorithm is applied to the cropped data (B). This algorithm discerns advantageous data points exhibiting optimal trade-offs, termed “knees” (curves with negative concavity) or occasionally “elbows” (curves with positive concavity).[8] Finally, a piecewise linear and cubic polynomial fit is employed to obtain the contact point as a fit parameter.

**Figure SI7:**
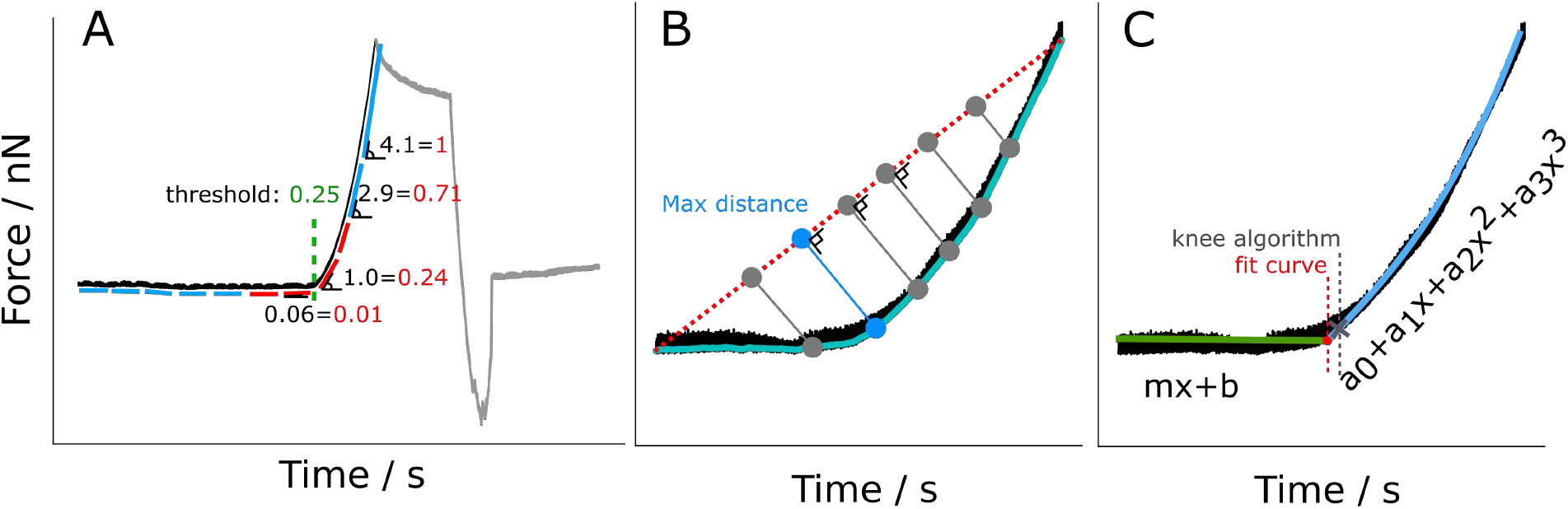
Algorithm to determine the precise contact point between the indenter and the curved epithelia in force curves.

### 3 Velocity dependence of mechanical parameters obtained from tension-based model - consistency of the model

Figure SI8 depicts how approach indentation velocity affects i) the slope at high strain, ii) the pre-stress *T*_0_, iii) the area compressibility modulus (scaling factor)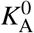, and iv) the fluidity represented by the power law exponent *β*. In this context, the viscoelastic model accurately captures how the velocity impacts the response. Higher approach velocities than 10 µm/s were not advisable due to hydrodynamic effects.[9] Lower velocities as shown here also turned out to be problematic as leakage of the lumen could overlay with viscoelastic relaxation processes as well as active responses of the cells subject to external stress.

**Figure SI8:**
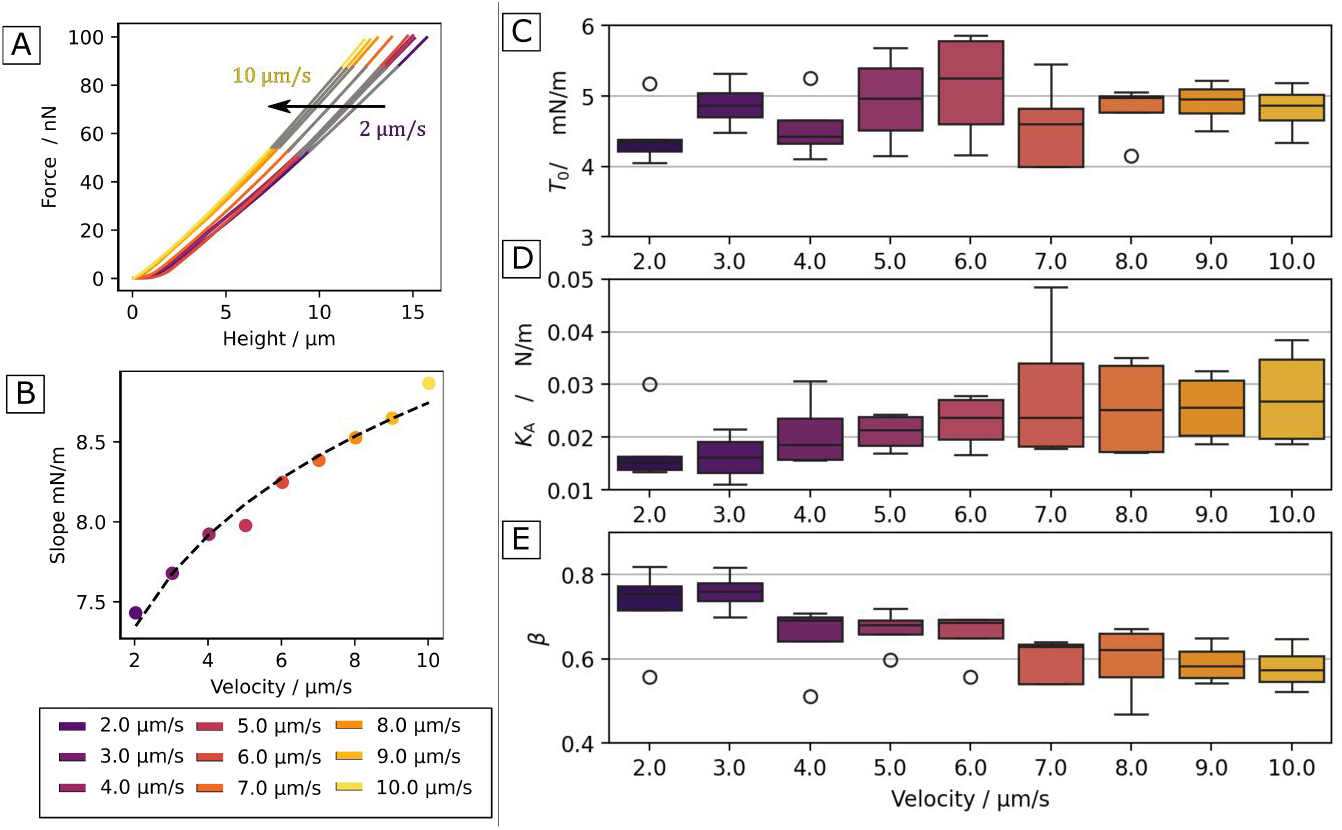
A) Average force curves each acquired at different indentation velocities. B) Slope of the indentation curve obtained from high strain as a function of loading rate. C-E) Mechanical parameters of the tension-based model remain predominantly unaffected by the approach velocity. Any slight rise in apparent stiffness could be ascribed to the elevated hydrodynamic drag acting on the indenter.

### 4 Pre-tension is consistent with different cantilever geometries

The tension of cysts were measured with two different cantilever geometries to show the outcome of the tension analysis is indenter-invariant. We selected a large spherical indenter and a flat surface (tipless cantilever) were employed, showing that there are no differences in tissue tension *T*_0_ regardless of the chosen setpoint.

**Figure SI9:**
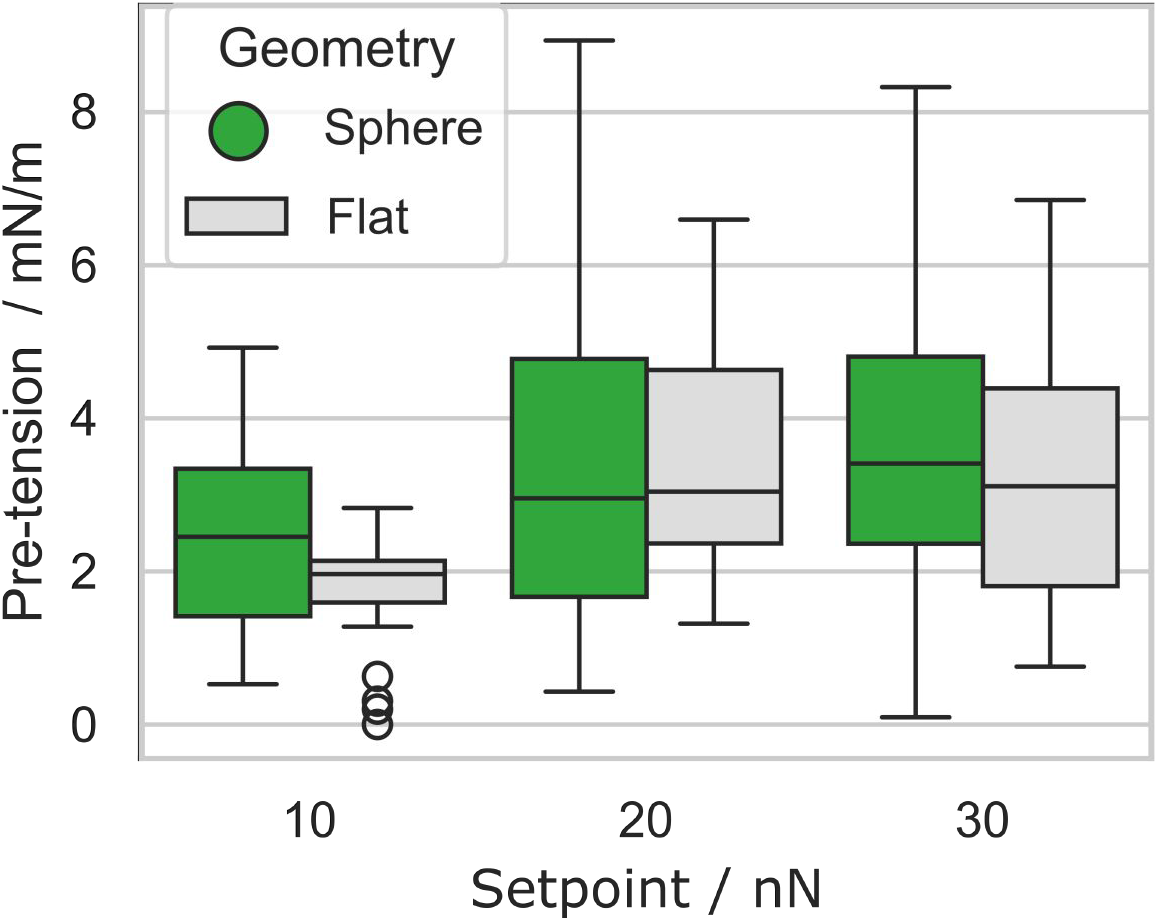
Tissue tension of cysts obtained from force curves using different cantilever geometries (parallel plate = flat and sphere *R* = 20 µm) and setpoints (maximal force).

### 5 Area dilatation of curved epithelia during indentation

Figure SI10 shows the increase in area dilatation *α* = Δ*A*/*A*_0_ as a function of indentation with a sphere (black) employing the tension model (see equation (36)). The area dilation remains well below 5% in the relevant indentation regime of this study supporting the validity of our restriction to the linear term in tension expansion:

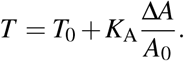

Indentation of the hemicysts generated even less areal strain compared to the cysts under typical experimental conditions such as size of the domes, indenter geometry, and indentation depth.

**Figure SI10:**
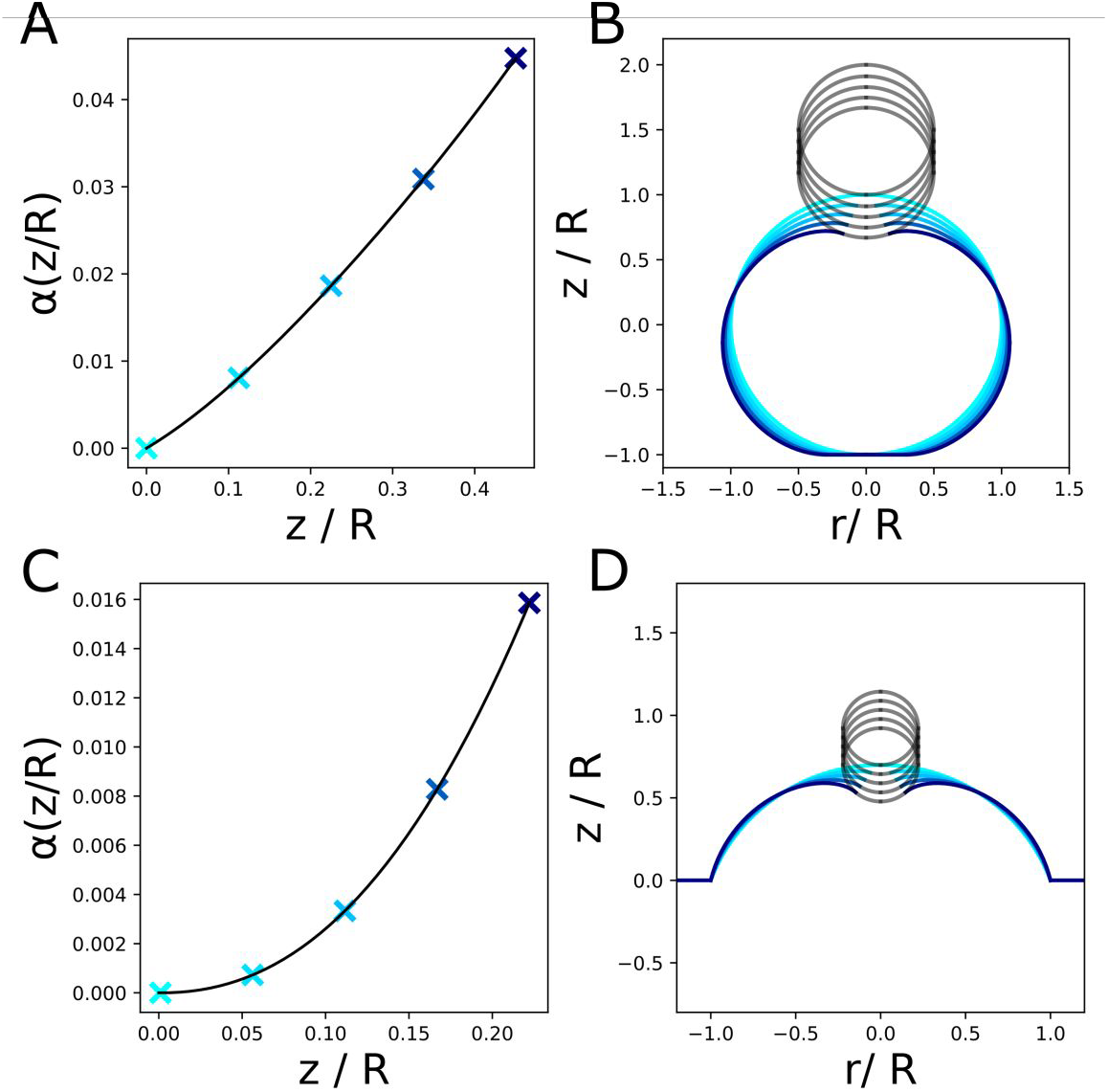
A/C) Area dilatation *α*(*z*/*R*) as a function of relative indentation depth *z*/*R*. Here, *R* represents the initial radius of the cyst (*R* = *R*_c_) or the base radius of the hemicyst (*R* = *R*_hc_), respectively. The accompanying illustrations of the curved epithelia during various indentation stages are displayed adjacent to the graphs (B/D).

### 6 Scaling of force curves

Scaling of experimental force data with different force setpoints ranging from 10 nN to 200 nN. This matches well the scaling regime with a slope in between 1 and 2 of the tension-based model shown in Fig. SI12.

**Figure SI11:**
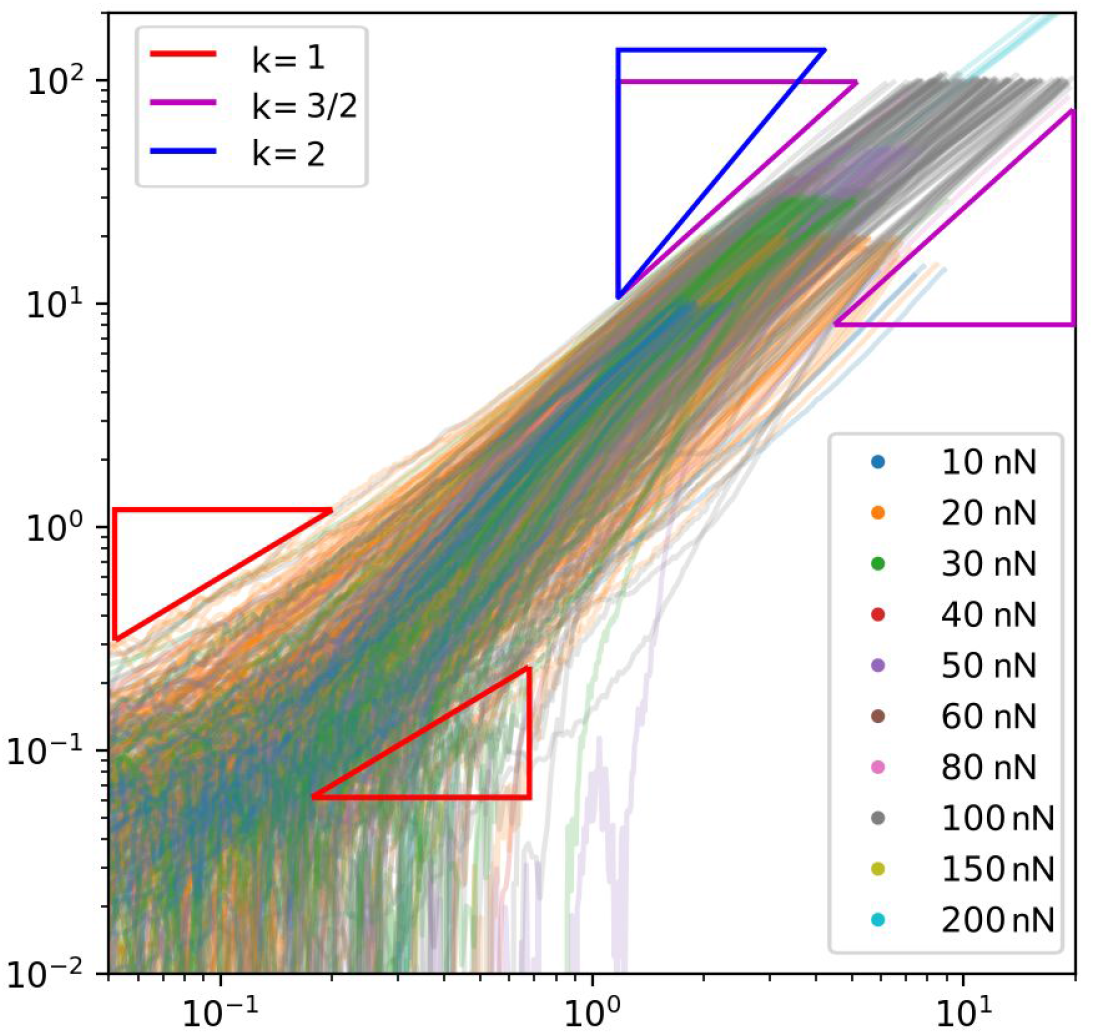
Force curves obtained from indentation of cysts shown in log-log plot to illustrate the scaling regimes. The plot comprises data from different setpoints (maximal forces, see legend).

### 7 Parameter space of the model -impact of mechanical parameters on the force curves

Figure SI12 illustrates the influence of pre-stress *T*_0_ (A/D), area compressibility modulus (scaling factor) 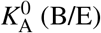, and fluidity (C/F) represented by the power law exponent *β* on force indentation and relaxation curves obtained from cysts deformed with a spherical indenter 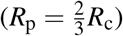. At low strain, the response is primarily elastic, influenced by the time-independent pre-stress (A/D), while at deeper indentation depths, 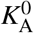 dominates (B/E). Increasing the area compressibility modulus affects both indentation and relaxation by increasing the dissipated energy. Higher *β* -values lead to a steeper slope during indentation and a faster decline of the yield force during relaxation (C/F).

**Figure SI12:**
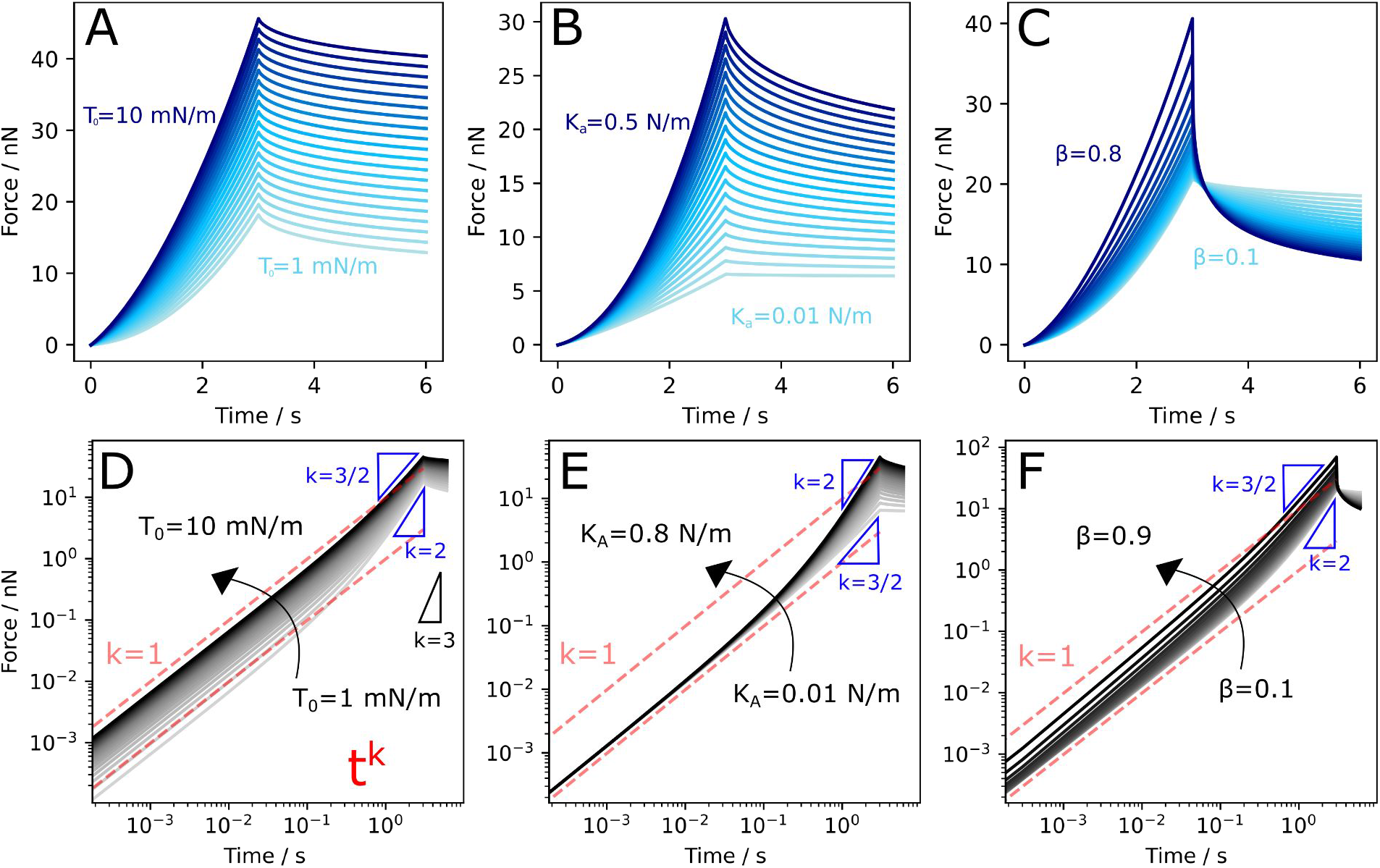
Simulations of indentation-relaxation curves. Impact of variations of pre-stress *T*_0_ (A), area compressibility 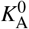(B) and fluidity *β* (C) on indentation relaxation curves as a function of time. A linear ramp with a constant indentation velocity (*ξ* = *vt*) is assumed. D-F) Log-log plot of (A-C) to illustrate the scaling *k* of the slope at various strains (see Fig. SI11).

### 8 Single cell geometry

The geometry of the apical cap of MDCK-II cells is approximated from CLSM measurements of a confluent monolayer.

**Figure SI13:**
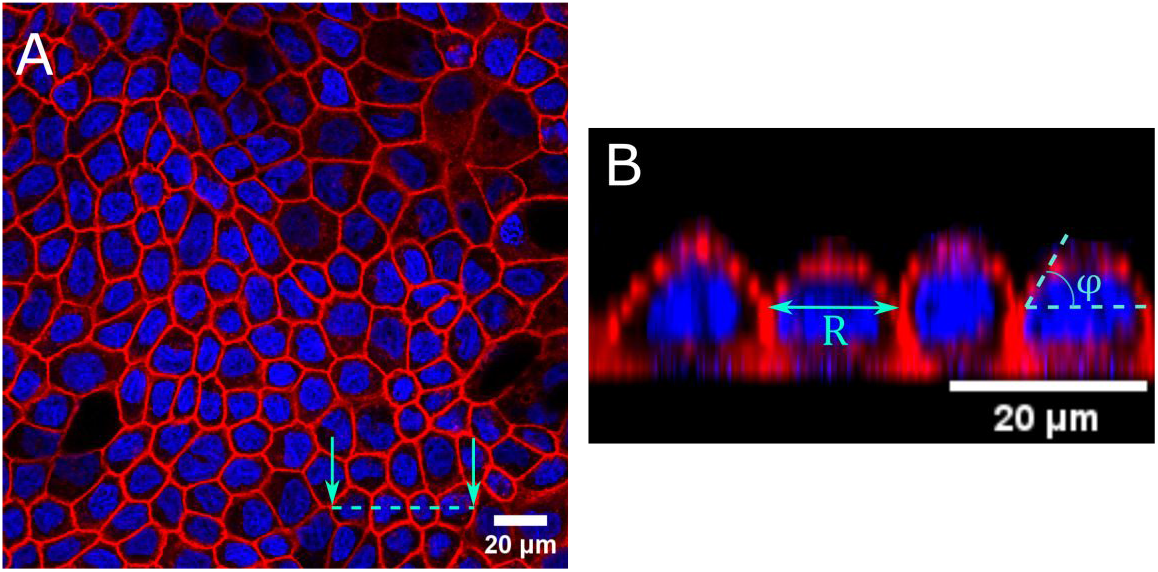
A) Confluent MDCK II cells on a regular culture dish. Cell membranes are stained with CellMask™ (red, 649 nm) and nuclei with Hoechst™ (blue, 405 nm). B) Cell geometry is extracted from *xz*/*yz*-cuts, allowing to measure the base radius and contact angle of the apical cap.

### 9 3D confocal images of cysts and hemicysts

The images in Fig. SI14 depict confocal images of cysts (A-D) and hemicysts (E-G), with the lumen stained through Dendra2-CD164 expression (fluorescent labeled mucin, green). Notably, there is a marked distinction in cell morphology of the tissue: cysts exhibit a more compact, cobblestone-like shape, whereas cells composing hemicysts are larger and less densely packed.

### 10 Scanning electron microscopy images of cysts and hemicysts

Figure SI15 shows two scanning electron microscopy (SEM) images depicting the contrasting polarity of cysts and hemicysts, respectively. The cyst exhibits no distinct surface features or cell boundaries, whereas the hemicyst reveals microvilli, with the apical side of the cells oriented outward.

**Figure SI14:**
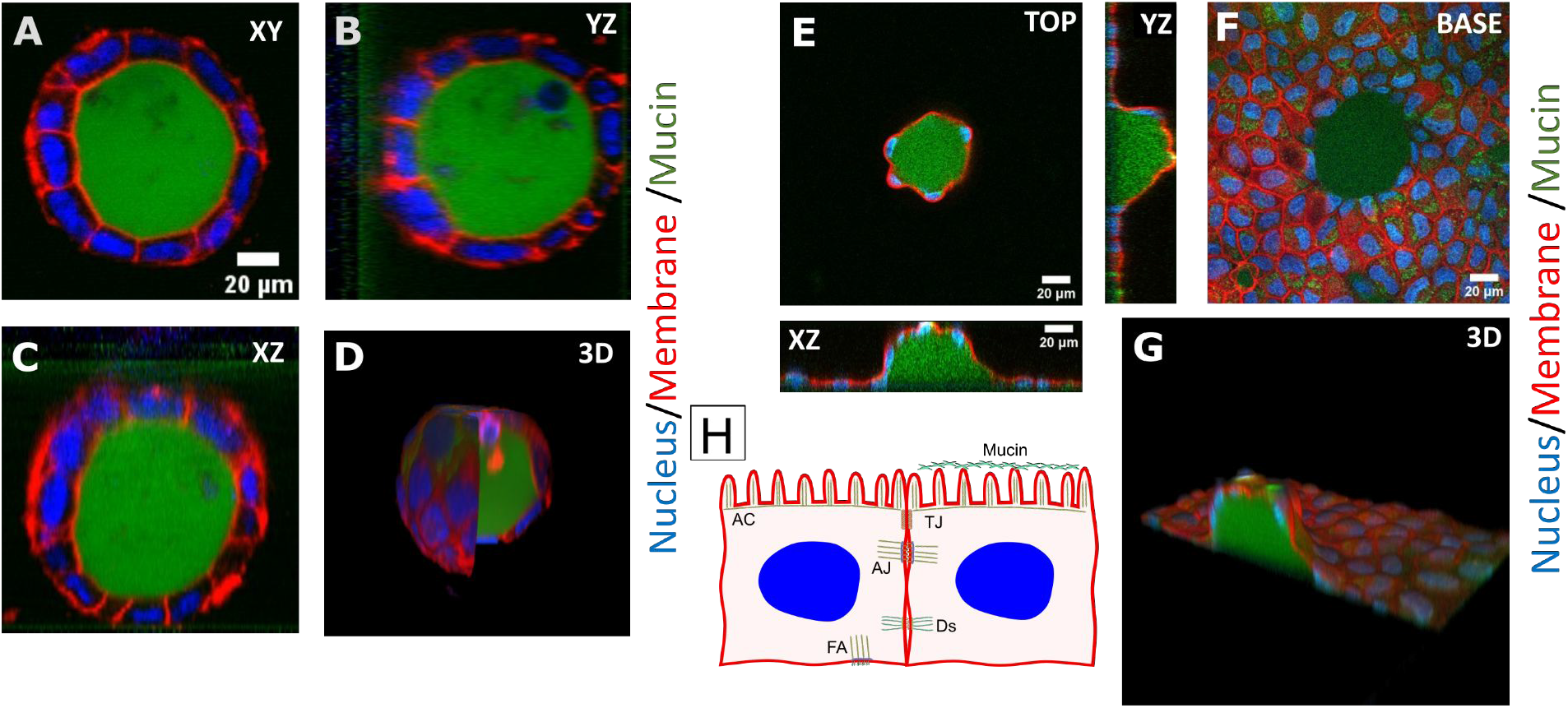
Confocal laser scanning microscopy images of cysts (left panels) and hemicysts (right panels) including a 3D reconstruction from *z*-scans. The nucleus is stained with DAPI (blue, 405 nm), the membrane with CellMask™ (red, 649 nm) and the lumen with the fluorescent, apically secreted sialomucin (Dendra2-CD164, green).

### 11 Nonlinear softening of tissue due to opening of cell-cell contacts

We simplify the individual bonds of an adherens junction attached to the cytoskeleton as springs arranged in parallel. Hence, a finite separation Δ*l* > 0 between the resting positions of bonds with tethers acting as springs (*k*_cad_) and the cytoskeleton acting as a transducer spring (*k*_cyt_) leads to an extension Δ*x*_cad_ of the tether spring and Δ*x*_cyt_ of the transducer spring with Δ*l* = Δ*x*_cad_ +Δ*x*_cyt_.[10] According to Hooke’s law, the corresponding forces to extend the springs are *F*_cad_ = *k*_cad_Δ*x*_cad_ and *F*_cyt_ = *k*_cyt_Δ*x*_cyt_. Assuming *n* closed bonds, the force balance is:

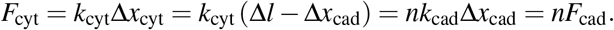

**Figure SI15:**
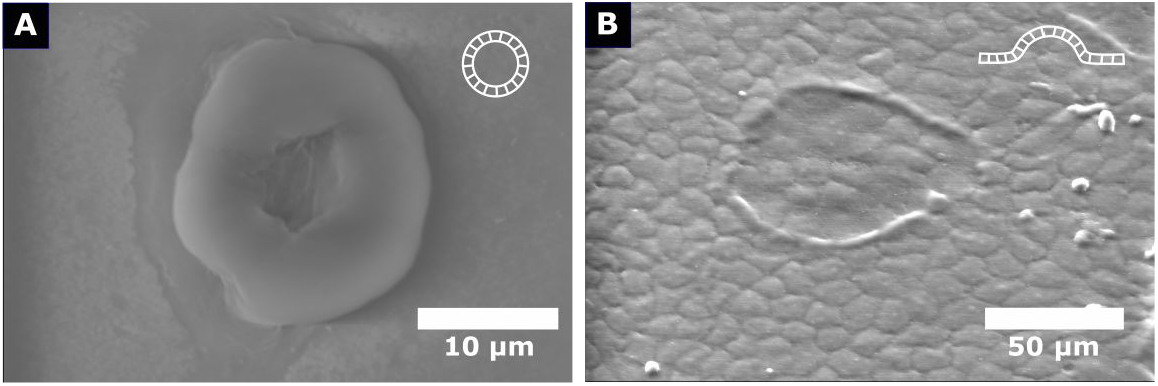
The scanning electron microscopy image displays the surface of a collapsed cyst (A) and a hemicyst (B). Notably, only the hemicyst’s surface reveals the presence of microvilli as the apical side faces outwards.

This result in the following expression for tether extension as function of *n*:

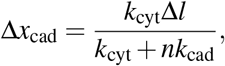

so that the force per bond is:

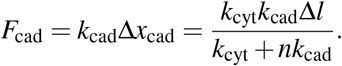

The extension of the spring representing the cytoskeleton is given by Δ*x*_cyt_ = Δ*l* − Δ*x*_cad_ so that

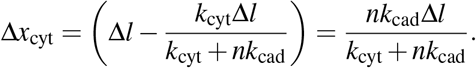

The resulting force acting on the cytoskeleton is

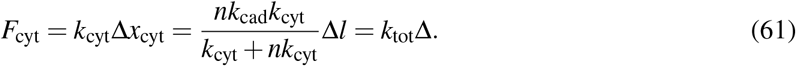

The effective Young’s modulus of the cyst’s shell can be envisioned as a linear combination of cells connected by springs that might open if tension rises:

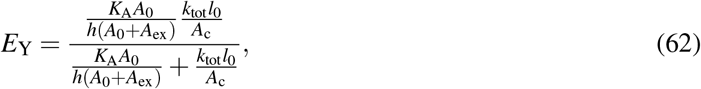

with *l*_0_ the resting length of the serial connection of springs, the contact area of the patch *A*_c_, and the effective spring constant 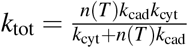, which is a function of the number of closed bonds *n*(*T*) depending on the tension *T*.

**Figure SI16:**
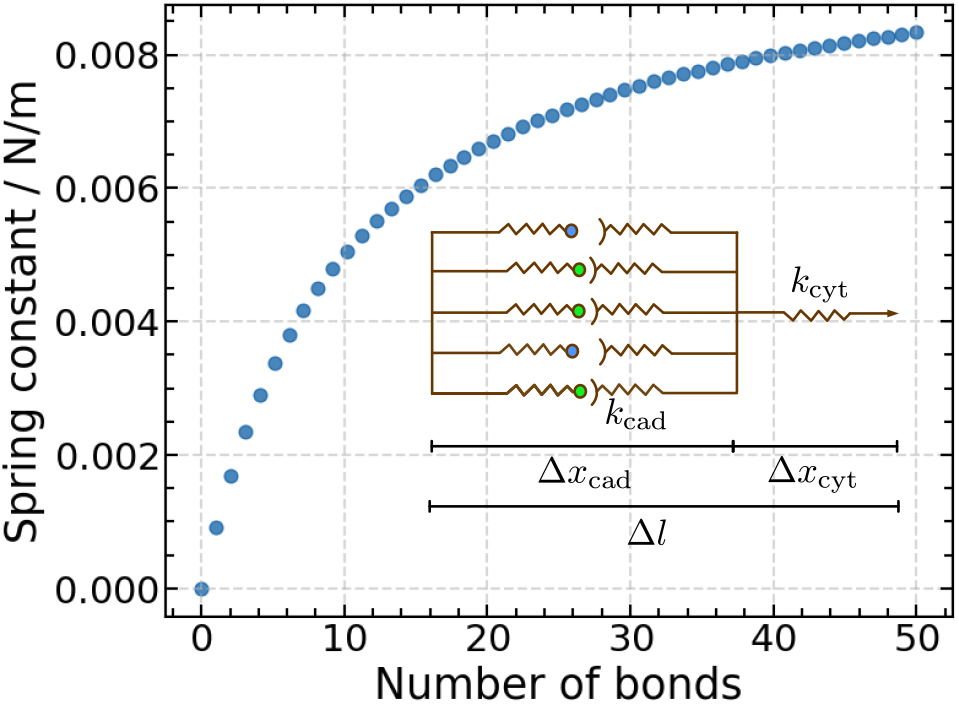
Effective spring constant *k*_tot_ as a function of closed bonds *n* (equation (61)) forming a patch of parallel tethers representing a single adherens junction (*k*_cad_ = 0.01 N/m, *k*_cyt_ = 0.1 N/m).

Bond number *n*(*T*) and excess interfacial area *A*_ex_ both effect the elastic modulus *E*_Y_ and by extending cell-cell-contacts and sacrificing surplus area mitigate the external stress and manage to maintain mechanical homeostasis in epithelial tissue.

**Figure SI17:**
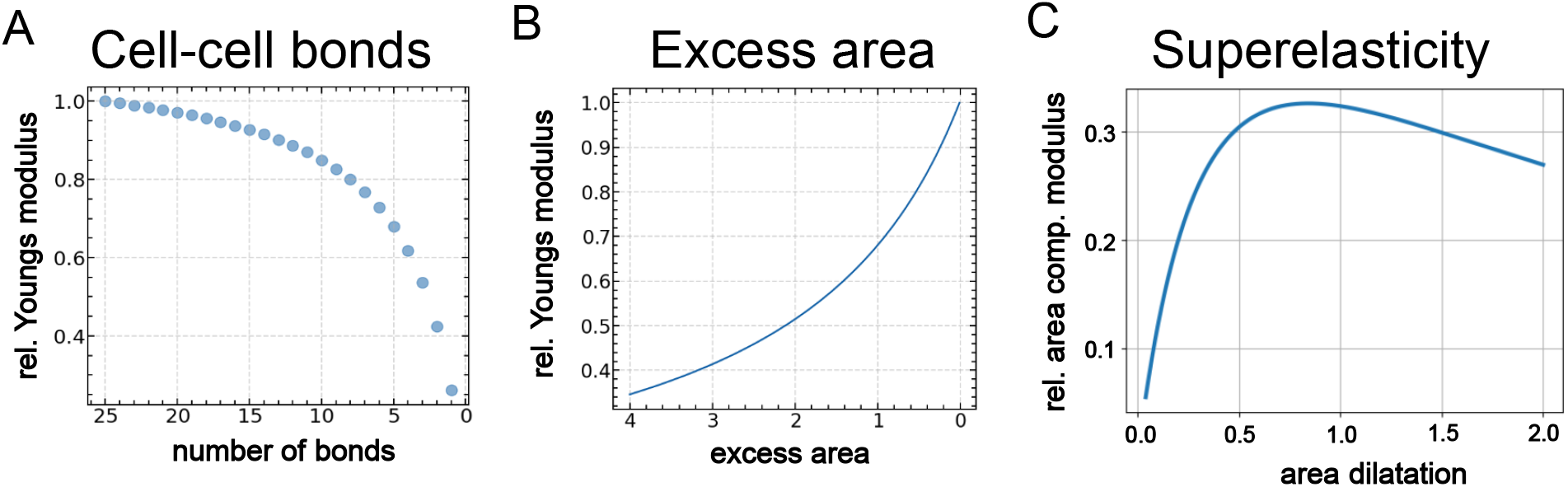
A/B) Change of *E*_Y_ (relative value, equation (62)) normalized to the initial value as a function of the number of bonds and relative excess area. While the loss of half the bonds between the cells only moderately affects the Young’s modulus (cooperative effect), the loss of excess area substantially increases the stiffness of the epithelia. C) Impact of area dilatation *α* on the area compressibility modulus illustrating the effect of superelasticity.

## Footnotes

1 Let the surface of separation undergo an infinitesimal displacement. At each point of the non-displaced surface we draw the normal. The length of the segment of the normal lying between the points where it intersects the displaced and non-displaced surface *δξ*. Then a volume element between the two surface is *δξ* d*A*, where d*A* is a surface element.

2 In the figure *δ A* = d*A*^*′*^ − d*A* = d*s*_1_d*s*_2_ (1 + *δξ* /*R*_1_ + *δξ* /*R*_2_) − d*s*_1_d*s*_2_

